# Niche contact during development programs hematopoietic stem cells

**DOI:** 10.64898/2026.11.17.767348

**Authors:** Nicole M. Woodhead, Sobhika Agarwala, Wantong Li, Bradley W. Blaser, Jianhong Ou, Rodolfo Calderon, Raquel Espin-Palazon, Clyde A. Campbell, Owen J. Tamplin

## Abstract

Fetal liver-derived hematopoietic stem and progenitor cells (HSPCs) have a higher capacity for self-renewal and reconstitution than bone marrow-derived HSPCs when transplanted into recipients. Therefore, stimulating self-renewal by reverting adult quiescent HSPCs to a fetal-like state could enable HSPC expansion for clinical applications. However, the gene regulatory networks and extrinsic signals involved in HSPC maturation that would need to be reversed are not well understood. Integrin alpha 4 (Itga4) has a conserved role for lodgment of HSPCs in the mammalian fetal liver, and the equivalent caudal hematopoietic tissue (CHT) in zebrafish. We hypothesized that comparison of wild-type HSPCs after lodgment in the CHT, and *itga4* mutant HSPCs that had bypassed the CHT, would reveal programming changes that occur during niche contact and maturation. We found *itga4* mutant HSPCs that did not lodge in the CHT are more immature, mitotic, myeloid-biased, and have an inflammatory signature. These features persisted into adulthood, suggesting that HSPCs kept in an immature state could result in chronic inflammation and reduced stem cell function.

## Introduction

Hematopoietic stem cells (HSCs) can self-renew and differentiate into any blood cell type. HSC contact with niche cells in the microenvironment is critical for development and regulation in the adult^1^. During development, in all studied vertebrates, HSCs emerge from the ventral wall of the dorsal aorta (DA) by budding from the hemogenic endothelium (HE)^2, 3^. In mammals, HSCs colonize the fetal liver (FL) where they undergo limited proliferation before migrating to the fetal bone marrow where they expand further before transitioning to quiescence^4–7^. Extensive work has been performed to define the differences between fetal and adult HSCs, including their distinct transcriptomes, proliferative status, and genome organization^8–20^. The gene regulatory networks of FL HSCs are considered the key to unlocking reversal of quiescence and stable expansion of adult HSCs for translational purposes^21, 22^. Furthermore, an understanding of the FL niche factors that regulate HSCs could lead to improvements in stable *ex vivo* expansion of HSCs^9, 11, 23–28^. Despite intense interest in the fetal-to-adult transition of HSCs, the intrinsic and extrinsic conditions required for HSC maturation are not fully understood.

The hematopoietic system is highly conserved in zebrafish, and its rapid development and transparent embryos allow easy tracking of hematopoietic stem and progenitor cells (HSPCs) in different sites of hematopoiesis^22, 29, 30^. HSPCs first emerge at ∼30 hours post fertilization (hpf) from the HE of the ventral DA^31, 32^. Multiple studies have shown that inflammatory signaling, including TNF-α, NF-κB, IFN-γ, IL6, IL1-β pathways, are involved in the endothelial-to-hematopoietic transition (EHT) in both zebrafish and mammals^33–43^. After emergence, HSPCs circulate and lodge in the FL-like caudal hematopoietic tissue (CHT) niche from 2-5 days post fertilization (dpf)^30, 44^. Evidence supports two types of HSPCs that colonize the mammalian FL and zebrafish CHT, embryo-restricted progenitors that sustain developmental hematopoiesis, and long-term definitive HSCs that persist into adulthood^5, 45–49^. Long-term definitive HSCs undergo quality control in the CHT niche to prevent poor clones from expanding and contributing to the adult HSC pool^50^. From the CHT, HSPCs then migrate to a perivascular niche in the anterior kidney region, where they remain throughout life and support adult hematopoiesis^29, 44^.

Intrinsic and extrinsic factors coordinate as HSPCs migrate between different sites in the embryo. One of these factors is integrin alpha 4 (Itga4) that heterodimerizes with integrin beta 1 (Itgb1) to form the VLA-4 protein complex on HSPCs^51^. The role of Itga4 on HSPCs, and its binding to Vcam1 on the niche, is highly conserved for HSPC lodgment in the mammalian FL and zebrafish CHT^50, 52–61^. An important distinction, however, is that failed FL hematopoiesis in mouse is embryonic lethal, while *itga4* loss-of-function zebrafish mutants survive because rapid development allows HSPCs to bypass the CHT and lodge directly in the larval kidney niche^52^. This feature of zebrafish *itga4* mutants provides an opportunity to compare intrinsic differences between wild-type (WT) HSPCs, and *itga4* mutant HSPCs that did not lodge in the CHT niche. We discovered that *itga4* mutant HSPCs had dysregulated inflammatory signaling pathways and increased proliferation rates. Importantly, *itga4* mutant HSPCs have an altered chromatin landscape, suggesting lodgment in the CHT niche is required for programmed epigenetic changes that contribute to HSPC maturation. Adult *itga4* mutant HSPCs have persistent inflammatory signaling, a distinct myeloid bias, and hyperproliferation that may result from altered accessibility of transcription factor (TF) binding sites and disrupted gene regulatory programs.

## Results

### HSPCs in *itga4* mutants have upregulated inflammatory pathways at 5 dpf

We confirmed that our previously established *itga4* mutant line (*itga4^UWT013^*)^62^, had decreased definitive HSPCs in the CHT at 5 dpf (Fig. S1), consistent with reduced HSPC retention and lodgment observed by us^63^ and others^52^. By 5 dpf HSPCs have already started to transition from the CHT to the kidney marrow niche^30, 44^. Therefore, Runx:mCherry+ HSPCs sorted from 5 dpf WT and *itga4* mutant larvae are developmentally homochronic but have either interacted with or bypassed the CHT niche, respectively. Bulk RNA-sequencing and analysis of these sorted populations (Fig. 1A), detected 335 differentially expressed genes (DEGs) (Fig. 1B, Table S1; False Discovery Rate (FDR)<0.05; 286 up- and 49 down-regulated transcripts in *itga4* mutant versus WT; minimum fold change (FC) of ±2 needed to identify significant transcripts). Gene Ontology (GO) and Gene Set Enrichment Analysis (GSEA) analysis of DEGs in *itga4* mutant HSPCs compared to WT found enrichment of immune response and inflammatory pathways (Fig. 1C-F). Oxidative phosphorylation and MYC target genes were underrepresented in *itga4* mutant HSPCs compared to WT (Fig. 1G,H). These results suggest Itga4-mediated interaction with the CHT niche is involved in regulating inflammatory and metabolic pathways that are known to fluctuate during HSPC development.

**Figure 1.**
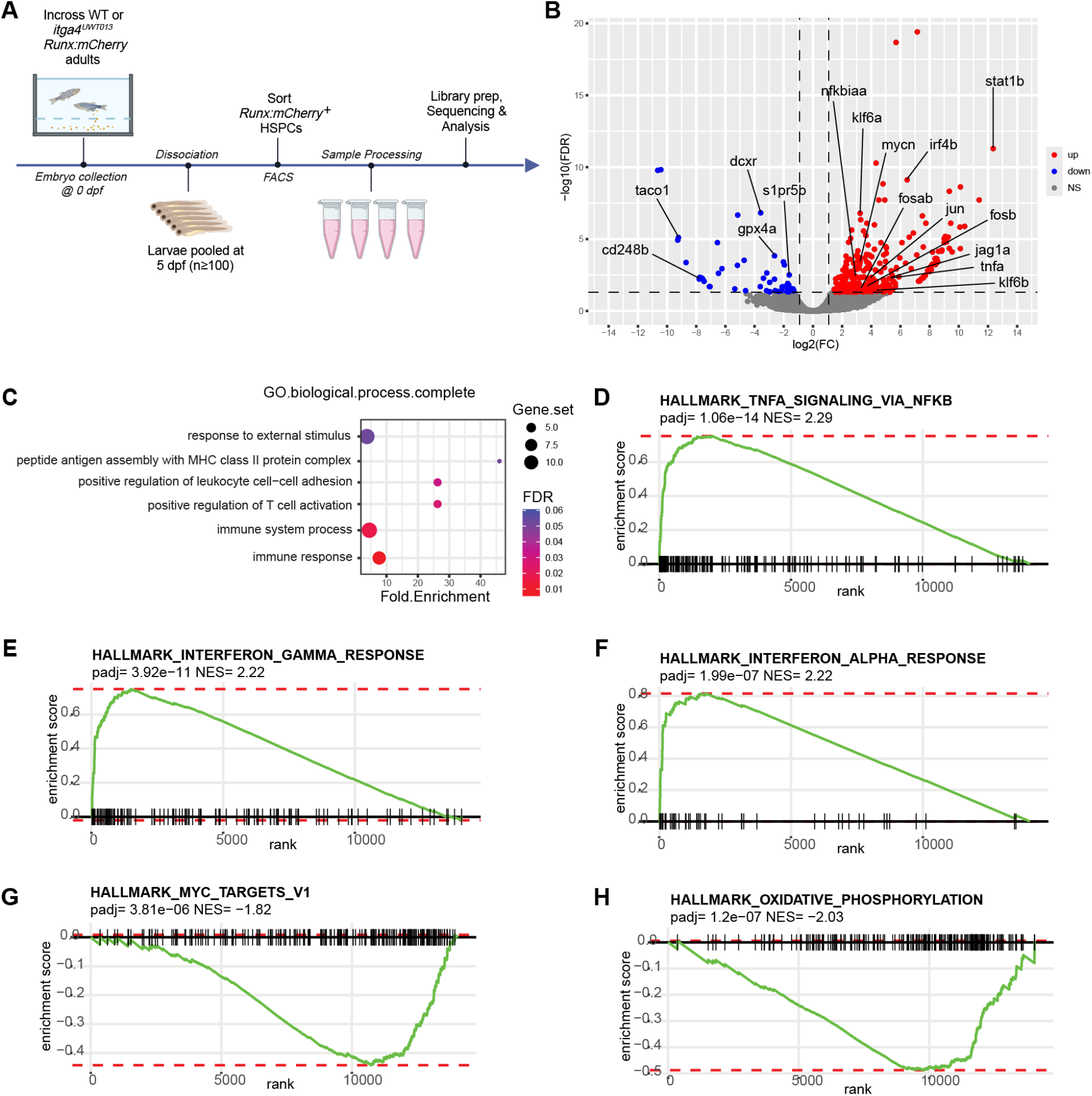
Bulk RNA-seq of Runx:mCherry+ HSPCs sorted from pooled *itga4^UWT013^*larvae reveal an inflammatory signature compared to WT at 5 dpf. **A)** Bulk RNA-seq workflow (created with Biorender). Experiment performed once and includes n=4 WT and n=3 *itga4^UWT013^*biological replicates, n≥100 pooled larvae per replicate. **B)** Volcano plot of differentially expressed genes (DEGs) between HSPCs from *itga4^UWT013^*and WT at 5 dpf. Red dots (right side) represent genes upregulated in *itga4^UWT013^*compared to WT, blue dots (left side) are genes downregulated in *itga4^UWT013^*compared to WT, grey indicates non-significant genes (vertical dashed lines indicate log2(FC)=±1, horizontal dashed line indicates FDR=0.05). **C)** Dotplot of gene ontology (GO) analysis from DEGs between *itga4^UWT013^* and WT HSPCs (panel **B**). Dot size indicates number of DEGs representing GO term, dot color represents significance value (red is low FDR, blue is high), fold enrichment describes *itga4^UWT013^*compared to WT HSPCs. **D-H)** Top Hallmark GSEA pathways represented by DEGs between *itga4^UWT013^* and WT HSPCs (panel **B**). Positive NES is upregulated in *itga4^UWT013^* compared to WT: **D)** TNFA_SIGNALING_VIA_NFκB; **E)** INTERFERON_GAMMA_RESPONSE; **F)** INTERFERON_ALPHA_RESPONSE. Negative NES is upregulated in WT compared to *itga4^UWT013^*; **G)** MYC_TARGETS_V1; **H)** OXIDATIVE_PHOSPHORYLATION. See also **Figure S1 and Table S1**.

### scRNA-seq reveals HSCs in *itga4* mutants at 5 dpf have an immature profile

We performed single cell RNA-seq (scRNA-seq) on sorted Runx:mCherry+ HSPCs from 5 dpf pooled WT and *itga4* mutant larvae. After batch correction and cluster annotation, we found that sorting Runx:mCherry+ HSPCs captures erythroid progenitors, common progenitors, definitive HSCs, neutrophils, and small populations of endothelial and stromal cells (Fig. 2A, S2A-C). As expected, based on our previous studies^30^, the highest level of *mCherry* transgene expression was in the HSC cluster, and lower levels were in more differentiated progenitors (Fig. 2A, S2D). To resolve the heterogeneity within the HSC cluster, WT- or *itga4* mutant-enriched neighborhoods were defined using differential abundance analysis and then characterized by pseudobulk differential gene expression analysis using embryo pools as the biological replicate unit^64^ (Fig. 2B,C, Table S2). We detected 565 transcripts enriched in the WT-enriched neighborhood and 467 transcripts enriched in the *itga4* mutant-enriched neighborhood (Benjamini-Hochberg (BH) adjusted p-value<0.05). This showed persistent upregulation of AP-1 factors and inflammatory pathway genes in the 5 dpf *itga4* mutant HSC neighborhood compared to WT (Fig. 2D). Consistent with the bulk RNA-seq analysis of 5 dpf *itga4* mutant Runx:mCherry+ HSPCs (Fig. 1D-F; Table S1), inflammatory cytokine genes were upregulated in the *itga4* mutant scRNA-seq HSC cluster (e.g., *nfκbiab* and *tnfaip3*; Fig. 2C,D; Table S2).

**Figure 2.**
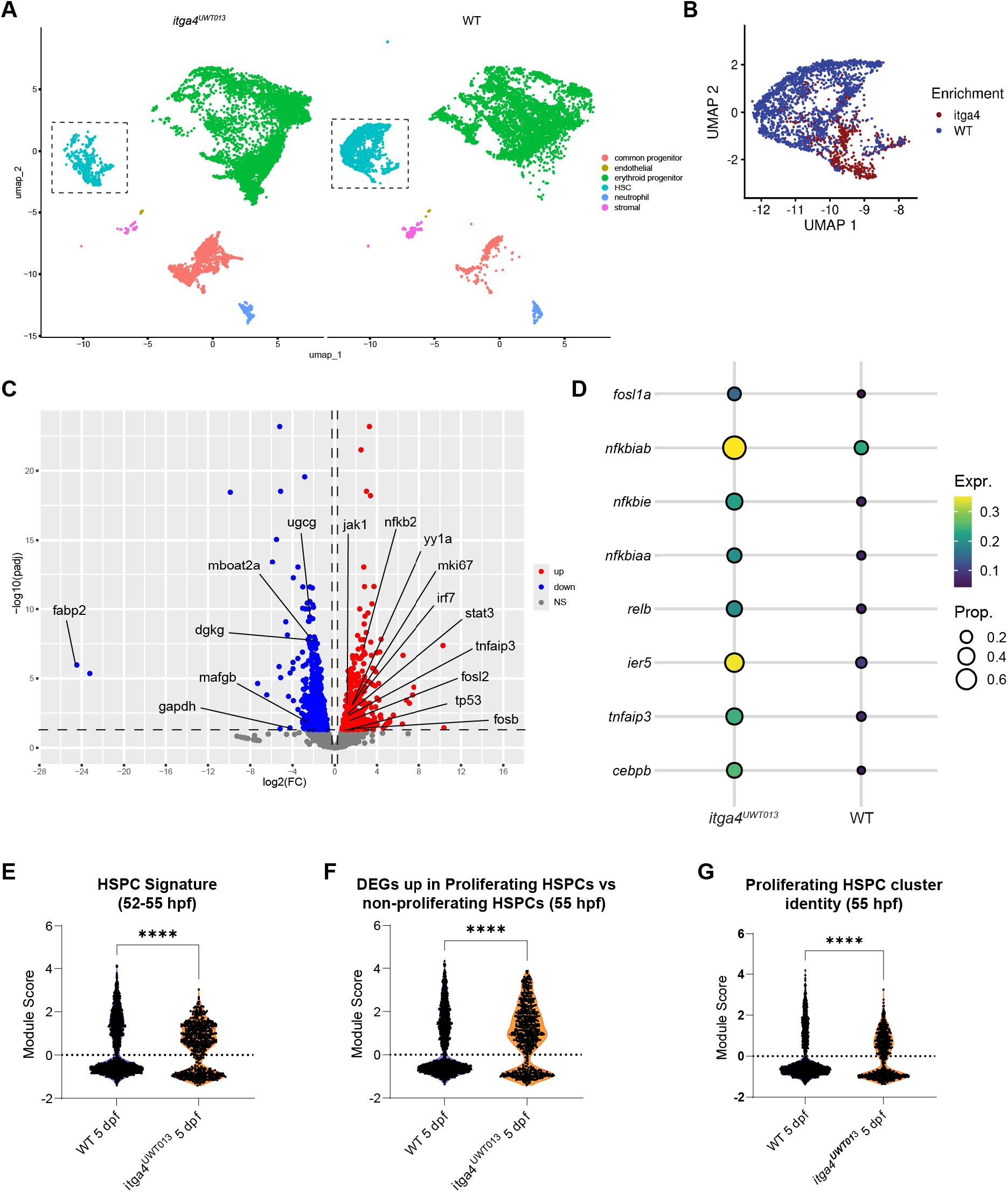
scRNA-seq of 5 dpf Runx:mCherry+ HSPCs sorted from pooled *itga4^UWT013^* larvae determine an HSC-intrinsic inflammatory signature compared to WT at 5 dpf. A) UMAP projection of *itga4^UWT013^* and WT samples. n=4 WT biological replicates with n≥100 pooled larvae per replicate from sorting to sequencing; n=3 *itga4^UWT013^* biological replicates with n≥100 pooled larvae per replicate were sorted. One *itga4^UWT013^* biological replicate was split into two technical replicates for sequencing. All samples were combined from three independent experiments and batch correction shown in **Fig. S2A**. Total cells from all replicates: n=12,687 WT and n= 13,967 *itga4^UWT013^*. Dashed boxes: HSC cluster used for further analysis (panels **B-G**). **B)** UMAP projection of HSC cluster (panel **A**) with cell classifications based on differential abundance “neighborhood” (red=*itga4^UWT013^*, blue=WT). **C)** Volcano plot of differentially expressed genes (DEGs) between *itga4^UWT013^* and WT (panel **B**). DEGs upregulated or downregulated in *itga4^UWT013^*vs WT on right (red dots) or left (blue dots), respectively. Non-significant genes (grey). Vertical dashed lines: log2(FC)=±0.25. Horizontal dashed line: BH adjusted p-value<0.05. **D)** Bubble plot of AP-1 factors and inflammatory pathway genes from HSC DEGs. **E-G)** Violin plots comparing module scores (MS) assigned to individual cells within the HSC cluster. MS positive (MS^+^) or negative (MS^-^) is higher or lower expression of module set genes vs control genes, respectively. Violin plot symbols represent single cells. Two-sided Chi-square analysis compared proportion of HSCs assigned to MS^+^ or MS^-^in 5 dpf WT vs *itga4^UWT013^*. Percent indicates proportion of positively scored HSCs. Degrees of freedom (df)=1. **E)** Chi-square=61.90, ****p<0.0001; MS^+^ 45.1% of WT (median=1.55, IQR=1.13) vs 62.9% of *itga4^UWT013^* (median=1.01, IQR=0.812); MS^-^54.8% of WT (median=-0.642, IQR=0.322) vs 37.1% of *itga4^UWT013^* (median=-0.935, IQR=0.277). **F)** Chi-square=139.5, ****p<0.0001; MS^+^ 44.0% of WT (median=1.65, IQR=1.36) vs 70.6% of *itga4^UWT013^* (median=1.41, IQR=1.34); MS^-^56.0% of WT (median=-0.636, IQR=0.317) vs 29.4% of *itga4^UWT013^*(median=-0.942, IQR=0.216). **G)** Chi-square=80.49, ****p<0.0001; MS^+^ 29.5% of WT (median=1.50, IQR=1.21) vs 48.8% of *itga4^UWT013^* (median=0.844, IQR=0.819); MS^-^70.4% of WT (median=-0.663, IQR=0.334) vs 51.2% of *itga4^UWT013^* (median=-0.953, IQR=0.243). See also **Figure S2 and Table S2**.

GSEA indicated that 5 dpf *itga4* mutant HSCs are enriched in TNFA signaling via NF-κB (Fig. S2E). Compared to the underrepresentation of MYC target genes in bulk RNA-seq of *itga4* mutant Runx:mCherry+ HSPCs (Fig. 1G) that includes HSCs and other progenitor populations (Fig. 2A), pseudobulk analysis of just the *itga4* mutant Runx:mCherry+ HSCs revealed a shift towards enrichment of MYC target genes (Fig. S2F). This is suggestive of a proliferative *itga4* mutant HSC population when analyzed without other progenitor populations. Consistently, we saw upregulation of proliferation-associated genes *mki67* and *yy1a* in *itga4* mutant HSCs compared to WT (Fig. 2C, Table S2). Inflammatory pathways are known to activate proliferative responses in HSCs shown by Ki67 staining^65–68^, and YY1 positively regulates HSC proliferation ^69,70^. Our results also show 5 dpf WT HSCs are enriched in PPAR signaling and lipid binding pathways compared to *itga4* mutant HSCs (Fig. S2G,H). Therefore, *itga4* mutant Runx:mCherry+ HSCs at 5 dpf have an inflammatory and proliferative state, while WT HSCs are undergoing a metabolic switch to the PPAR-fatty acid oxidation (FAO) pathway that promotes HSC maintenance^71^.

The inflammatory signature of *itga4* mutant HSCs at 5 dpf was reminiscent of newly formed HSPCs that activate TNF-α, NF-κB, IFN-γ, and IL1-β signaling during emergence from the DA^33–37^. We performed a module score analysis to compare 5 dpf WT and *itga4* mutant HSCs against published scRNA-seq data of early-stage 52-55 hpf WT HSPCs^26^. A significantly larger proportion of 5 dpf *itga4* mutant HSCs ranked positively with newly born 52-55 hpf HSPCs compared to WT 5 dpf HSCs (Fig. 2E). In addition, significantly more 5 dpf *itga4* mutant HSCs, compared to WT, ranked positively with genes related to HSPC proliferation (Fig. 2F), and the cluster identity of proliferating HSPCs at 55 hpf (Fig. 2G). These comparisons show 5 dpf *itga4* mutant HSCs that have bypassed the CHT maintain a more immature, inflammatory, and proliferative profile.

### *itga4* mutant HSPCs have an altered chromatin landscape at 5 dpf

We considered that not only the gene expression profile but also the chromatin landscape of *itga4* mutant HSPCs may differ from WT because of failed lodgment in the CHT. We sorted Runx:mCherry+ HSPCs from 5 dpf WT and *itga4* mutant larvae for bulk ATAC-sequencing and analysis. We detected 161 accessible regions of chromatin (“peak” signals) unique to WT HSPCs and 837 peaks unique to *itga4* mutant HSPCs (FC>±1.1, p<0.05), suggesting that epigenetic remodeling occurs in HSPCs after CHT niche interactions (Fig. 3A). Genome-wide distribution of accessible chromatin regions between WT and *itga4* mutant HSPCs, such as promoters, introns, and intergenic regions, remained similar (Fig. S3A). We hypothesized that HSPCs depend on contact with the CHT niche to undergo chromatin remodeling and initiate maturation programs. Regions with greater accessibility in *itga4* mutant HSPCs at 5 dpf, compared to WT, may have been open in early-stage immature HSPCs, and then remained open in the absence of CHT lodgment (Fig. 3A). This includes putative long-range enhancers for *klf6a* and *klf6b* genes that encode Krüppel-like TFs known to regulate hematopoietic and vascular development^72–74^ (Fig. 3B; Table S3). Gene loci involved in inflammatory pathways are also accessible in *itga4* mutant HSPCs at 5 dpf (Fig. 3B); these include *il6* that is involved in EHT^43^, *tnfsf18* that is implicated in fetal myelopoiesis^75^, and *nfkbie* that regulates HSC quiescence^76^. A parity plot confirmed that positive FC of chromatin accessibility at gene loci associated with hematopoietic TFs and inflammatory pathways (e.g., *klf6a*, *klf6b*, *nfkbie*) correlated with higher transcript expression levels (Fig. S3B). GO analysis of genes with peaks that are uniquely accessible in *itga4* mutant HSPCs compared to WT are indicative of HSPC activation, as GO terms relate to migration, increased metabolism, and ribosome assembly (Fig. S3C; Table S3). Examples of differentially accessible chromatin peaks in *itga4* mutant HSPCs are found near important hematopoietic specification TFs that are active in early emergent HSPCs^14, 77^ (e.g., *meis1a*, *ets1*, *gata2a*, *egr1*; Fig. S4A-D, Table S3). Inflammatory factor *tnfb* also had increased accessibility in *itga4* mutant HSPC upstream promoter regions (Fig. S4E). Interestingly, a gene that was more accessible in WT HSPCs was the TF *hoxb4* that is a potent driver of HSC self-renewal^78^ (Fig. S4F). Together, these data show that WT HSPCs that lodged in the CHT, or *itga4* mutant HSPCs that did not, have significantly different epigenetic programming.

**Figure 3.**
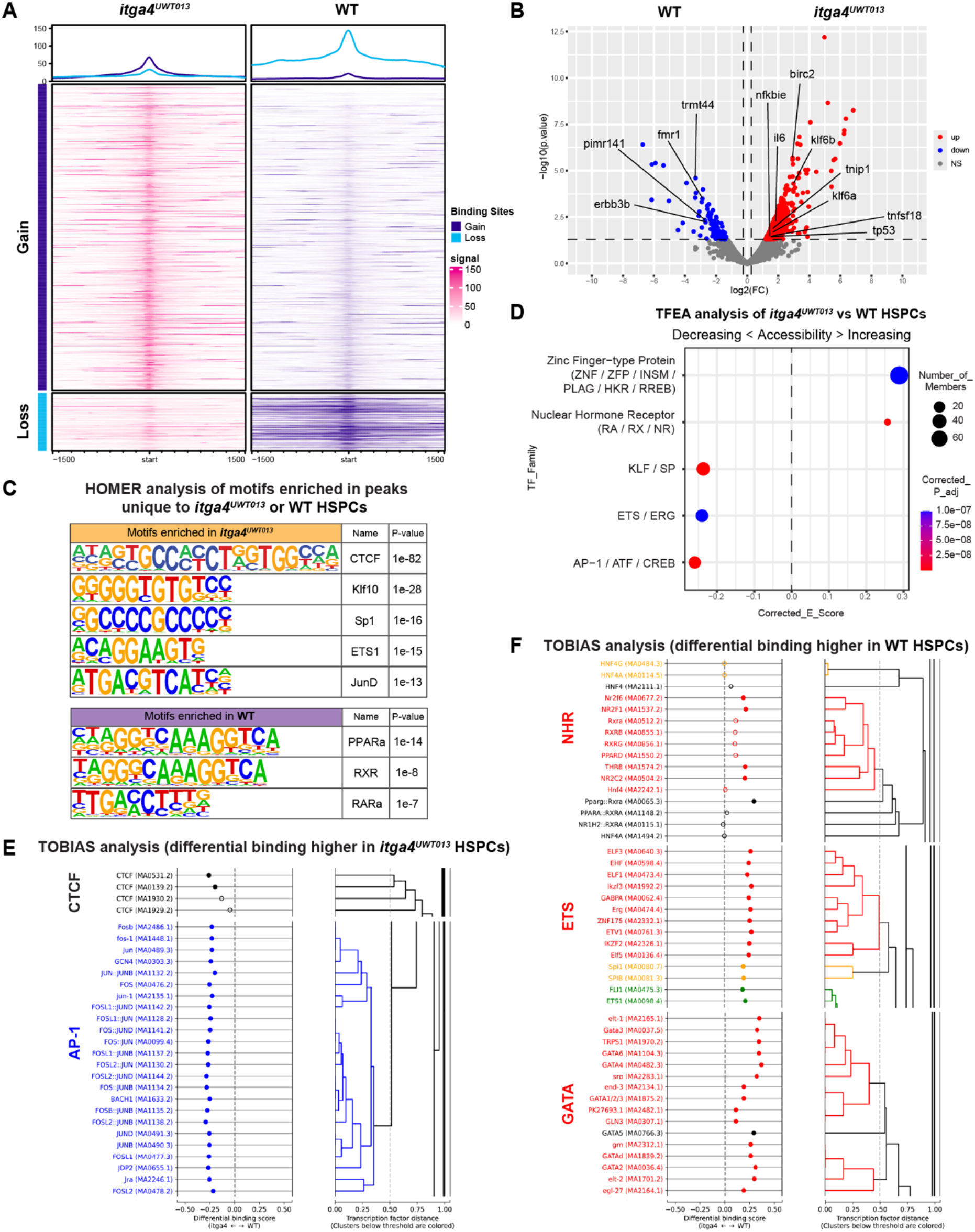
Bulk ATAC-seq of Runx:mCherry+ HSPCs sorted from pooled *itga4^UWT013^* larvae reveal differences in the chromatin landscape compared to WT at 5 dpf. **A)** Metaplots (top) and heatmaps (bottom) showing differential chromatin accessibility between *itga4^UWT013^* and WT HSPCs (plots centered on transcription start site (TSS); n=3 WT and n=3 *itga4^UWT013^*biological replicates; n≥100 pooled larvae per replicate. “Gain” and “Loss” describes increased or decreased accessibility, respectively, in chromatin regions from *itga4^UWT013^* (left) vs WT (right). Upper: Metaplot average accessibility signal (purple “gain” curves and blue “loss” curves). Lower: Heatmap accessibility signal from all peaks (stronger color saturation=greater accessibility; left/pink=*itga4^UWT013^*; right/purple=WT). **B)** Volcano plot of gene loci with unique accessibility in *itga4^UWT013^* vs WT HSPCs (mapped from “gain” and “loss” peaks; panel **A**). Red dots (right) represent upregulated accessibility in *itga4^UWT013^* vs WT; blue dots (left) represent downregulated accessibility in *itga4^UWT013^* vs WT; grey indicates no significance; vertical dashed lines indicate log2(FC)=±0.25; horizontal dashed line indicates p=0.05. **C)** Top predicted TF families from HOMER analysis of unique peaks. Binding motif, name, and p-value represents top result in family. **D)** Bubble plot summarizing predicted TFEA activity of TF families across *itga4^UWT013^* vs WT HSPC epigenomic landscapes (panels **A-B**). Dot size indicates number of TFs represented by family (corrected p adj<0.05). Families containing <4 TFs not shown. Corrected enrichment score (E Score) and p adj. assigned from highest-ranking TF in the family. Between *itga4* mutant and WT HSPCs, TFEA predicted regions with decreasing accessibility (negative E score) and increasing accessibility (positive E score). **E-F)** Results from TOBIAS TF binding prediction analysis comparing *itga4^UWT013^* and WT HSPCs. Hierarchical clustering based on motif similarity (color=similar; black=unique). **E)** TFs predicted to gain binding sites in *itga4^UWT013^* vs WT; grouped families represent CTCF and AP-1 factors. **F)** TFs predicted to gain binding sites in WT vs *itga4^UWT013^*; grouped families represent GATA, ETS and nuclear hormone receptor (NHR) factors. See also **Figure S3-4 and Table S3**.

To elucidate the regulatory programs that are active in WT or *itga4* mutant HSPCs at 5 dpf, we performed HOMER analysis^79^ to identify motifs enriched in regions of accessible chromatin (Fig. 3C, Table S3). We hypothesized that peaks unique to *itga4* mutant HSPCs, and remain open at 5 dpf, would normally lose accessibility and close after contact with the CHT niche. We found peaks unique to *itga4* mutant HSPCs were enriched in CTCF, KLF, SP1, ETS, and AP-1 TF motifs (Fig. 3C; Table S3). This is consistent with significantly higher expression of KLF and AP-1 TF genes in 5 dpf *itga4* mutant HSPCs, shown by bulk RNA-seq (Fig. 1B) and scRNA-seq (Fig. 2C,D). Peaks unique to 5 dpf WT HSPCs were enriched in nuclear hormone receptors (NHRs; e.g., PPARa) and retinoic acid (RA) receptor motifs (e.g., RARa and RXR) (Fig. 3C; Table S3). PPAR signaling genes were also enriched in WT HSCs at 5 dpf (Fig. S2G). PPAR-FAO pathway promotes HSC maintenance^71^, and RA signaling is associated with HSC dormancy^80^, suggesting WT HSPCs at 5 dpf are undergoing maturation, while *itga4* mutant HSPCs remain activated.

Next, we used TF enrichment analysis (TFEA)^81^ and TOBIAS^82^ analyses to predict changing TF activity and footprints, respectively, between the WT and *itga4* mutant states. At 5 dpf *itga4* mutant HSPCs exhibit characteristics of more immature, newly emerged HSPCs (∼52-55 dpf; Fig. 2E-G). Therefore, we hypothesized that comparison of 5 dpf *itga4* mutant and WT HSPCs would reveal the mechanisms of transition between immature and mature states, respectively. Accordingly, TFEA predicted decreasing accessibility of regions containing KLF, ETS, and AP-1 family motifs, and increasing accessibility of zinc finger and NHR motifs between *itga4* mutant and WT HSPCs (Fig. 3D, Table S3), consistent with HOMER analysis (Fig. 3C). TOBIAS normalizes accessibility signals and controls for Tn5 transposase insertion bias when predicting TF activity across the chromatin landscape^82^. Like HOMER and TFEA, TOBIAS predicted greater binding of AP-1 and CTCF factors in *itga4* mutant HSPCs, and NHR TFs in WT HSPCs (Fig. 3E,F, Table S3). TOBIAS also identified greater binding of GATA factors in WT HSPCs. Contrary to HOMER and TFEA, TOBIAS predicted ETS motifs as more actively bound in WT HSPCs than *itga4* mutant HSPCs at 5 dpf (Fig. 3F, Table S3). The different predicted results for ETS binding may reflect that HSPCs are continuously changing as they transition between states^80^. The AP-1 factor regulatory programs consistently detected in *itga4* mutants at 5 dpf are associated with early and emergent HSPCs^83, 84^, whereas RA programs in WT HSPCs indicate a transition to a more quiescent state^41, 80^.

### *itga4* mutant HSPCs show a higher *in vivo* mitotic rate and NF-κB activity

The gene expression profile and epigenetic landscape of *itga4* mutant HSPCs at 5 dpf suggests a more active and inflammatory state. Therefore, we decided to assess *in vivo* mitotic activity using the transgenic reporter line *zFUCCI* to mark cells in G_2_/S/M phases with monomeric Azami Green fluorescent protein^85^. Together with *Runx:mCherry* to mark HSPCs^30^, we imaged co-expression with *zFUCCI* in the CHT at 3 dpf (Fig. 4A). Although the absolute number of HSPCs in *itga4* mutants is lower, and their lodgment is transient^52, 63^, we could still determine the cell cycle phase of the remaining HSPCs (Fig. 4A,B). Analysis was performed by three-dimensional rendering of cell surfaces (Imaris software), and at least 95% overlap of zFUCCI and mCherry in HSPCs to designate co-expression (Fig. 4A, S5). Although the absolute number of HSPCs was significantly lower in *itga4* mutants compared to WT, there was a much higher proportion of HSPCs undergoing division (Fig. 4B). This *in vivo* data is consistent with upregulation of cell cycle regulators in *itga4* mutant HSPCs based on scRNA-seq (Fig. 2C) and chromatin accessible genes (Fig. 3B).

**Figure 4.**
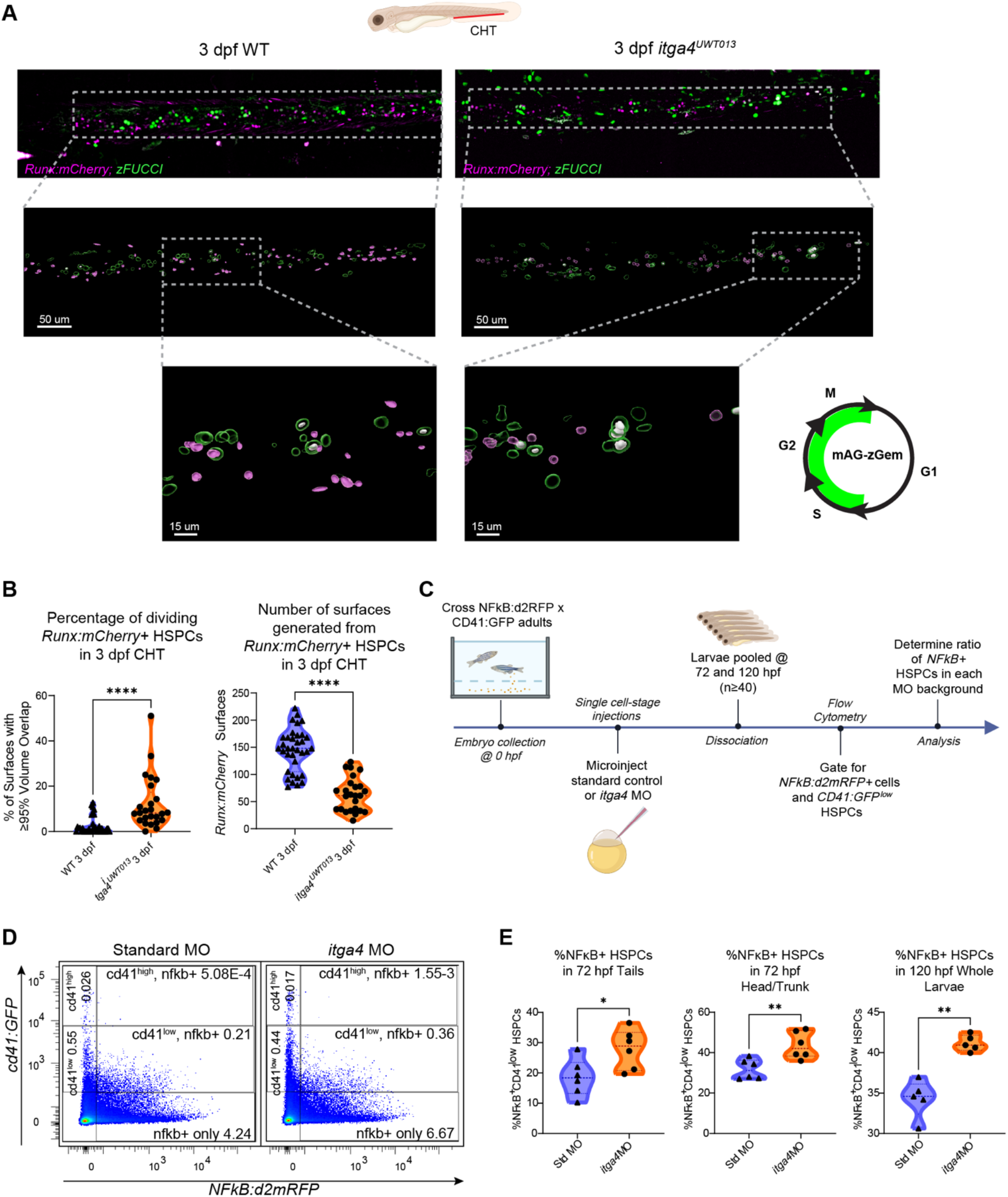
Cell division and NF-κB signal is increased in *itga4* mutant HSPCs. **A)** Transgenic *Runx:mCherry;zFUCCI* larvae were imaged at 3 dpf from WT (n=35 biological replicates, one larva/replicate, representative on left) and *itga4^UWT013^* (n=25 biological replicates, one larva/replicate, representative on right). Two independent experiments with replicates combined for analysis. Larva diagram (top) indicates orientation with dorsal up and anterior left (created with Biorender). Images were processed and analyzed via surface rendering and quantification of volume overlap as a proxy for co-expression of fluorescent signal (see **Methods**). Boxes indicate region zoomed below. Surfaces colors: green=zFUCCI; magenta=Runx:mCherry; white=overlap ≥95%. **B)** Proportion of zFUCCI+/Runx:mCherry+ cells among total Runx:mCherry+ cells (i.e., white/magenta surfaces) in the CHT. Purple violin with triangle points, left, represents WT (median=1.176% mitotic HSPCs, IQR=2.326%). Orange violin with circle points, right, represents *itga4^UWT013^* (median=8.197% mitotic HSPCs, IQR=12.33%). Analysis via unpaired Mann-Whitney test (****p<0.0001). Total number of magenta surfaces was significantly lower in *itga4^UWT013^*vs WT (median= 61 and IQR=46.5 vs median= 148 and IQR= 63; unpaired Mann-Whitney test, ****p<0.0001). **C)** Workflow for assessing NF-κB activity in HSPCs during CHT stage via flow cytometry (created with Biorender). Standard control or *itga4* MO injected at single cell stage. Larvae at 72 or 120 hpf pooled and dissociated for flow cytometry analysis. Determine proportion of HSPCs with active NF-κB. **D)** Representative gating strategy for flow cytometry: Standard control MO (left); *itga4* MO (right); y-axis cd41:GFP; x-axis NFκB:d2mRFP. cd41:GFP^low^/ NF-κB+ gate represents HSPCs with active NF-κB. **E)** Left: Percentage of HSPCs in tails with active NF-κB at 72 hpf (Control MO median=18.4 and IQR=10.2; *itga4* MO median=28.9 and IQR=12.6; *p=0.0411). Center: Percentage of HSPCs in trunks with active NF-κB at 72 hpf (Control MO median=31.2 and IQR=8.97; *itga4* MO median=42.1 and IQR=12.7; **p=0.0043). Right: Percentage of HSPCs in whole larvae with active NF-κB at 120 hpf (Control MO median=34.6 and IQR=3.72; *itga4* MO median=40.9 and IQR=1.86; **p=0.0079). Analysis via unpaired Mann-Whitney test. See also **Figure S5-6 and Methods**.

Pro-inflammatory signals, including NF-κB signaling, are required for HSPC emergence from the DA^34, 35^. Our genomic data suggests *itga4* mutant HSPCs at 5 dpf retain an immature and pro-inflammatory profile. We wanted to determine if NF-κB signaling activity was elevated *in vivo* in 5 dpf *itga4* knock-down HSPCs. We used the destabilized RFP transgenic reporter line *NF-κB:d2mRFP*^35^, together with *cd41:GFP*^86^, to detect active NF-κB signaling *in vivo* in HSPCs. Single-cell embryos were injected with either control morpholino (MO) or previously validated *itga4* knockdown MO^52^, then dissociated at 72 or 120 hpf for flow cytometry (Fig. 4C, S6). We dissected tails from head and trunk at 72 hpf to determine if there were regional differences in the distribution of cd41:GFP+ cells. However, our results were consistent with whole larvae at 120 hpf—independent of stage or region, we always found a consistent increase in the proportion of NF-κB+;cd41:GFP+ HSPCs after *itga4* MO knockdown compared to controls (Fig. 4D). Together, using *in vivo* reporters of cell cycle and NF-κB activity, we confirmed the higher mitotic rate and inflammatory pathway activity predicted by our genomic analyses.

### Adult *itga4* mutant HSPCs maintain an inflammatory and proliferative profile

To understand how *itga4* loss of function impacts HSPCs beyond development and into adulthood, we compared whole kidney marrow (WKM) from young 6 months post fertilization (mpf) WT, *itga4* mutant, and *vcam1b* mutant (*vcam1b^UWT007^*)^62^ (Fig. 5A). We included adult *vcam1b* mutants to determine if their HSPCs had a similar phenotype to *itga4* mutants, due to disruption of Itga4-Vcam1 HSPC-niche adhesion. Alternatively, HSPCs in *itga4* mutants may differ from those in *vcam1b* mutants because of the additional contribution of Itga4-dependent “outside-in” signaling^87, 88^. We also included aged WT adults (24 mpf) to determine if there were similarities between *itga4* mutant marrow and the chronic inflammation present in aged marrow^89^ (Fig. 5A).

**Fig 5.**
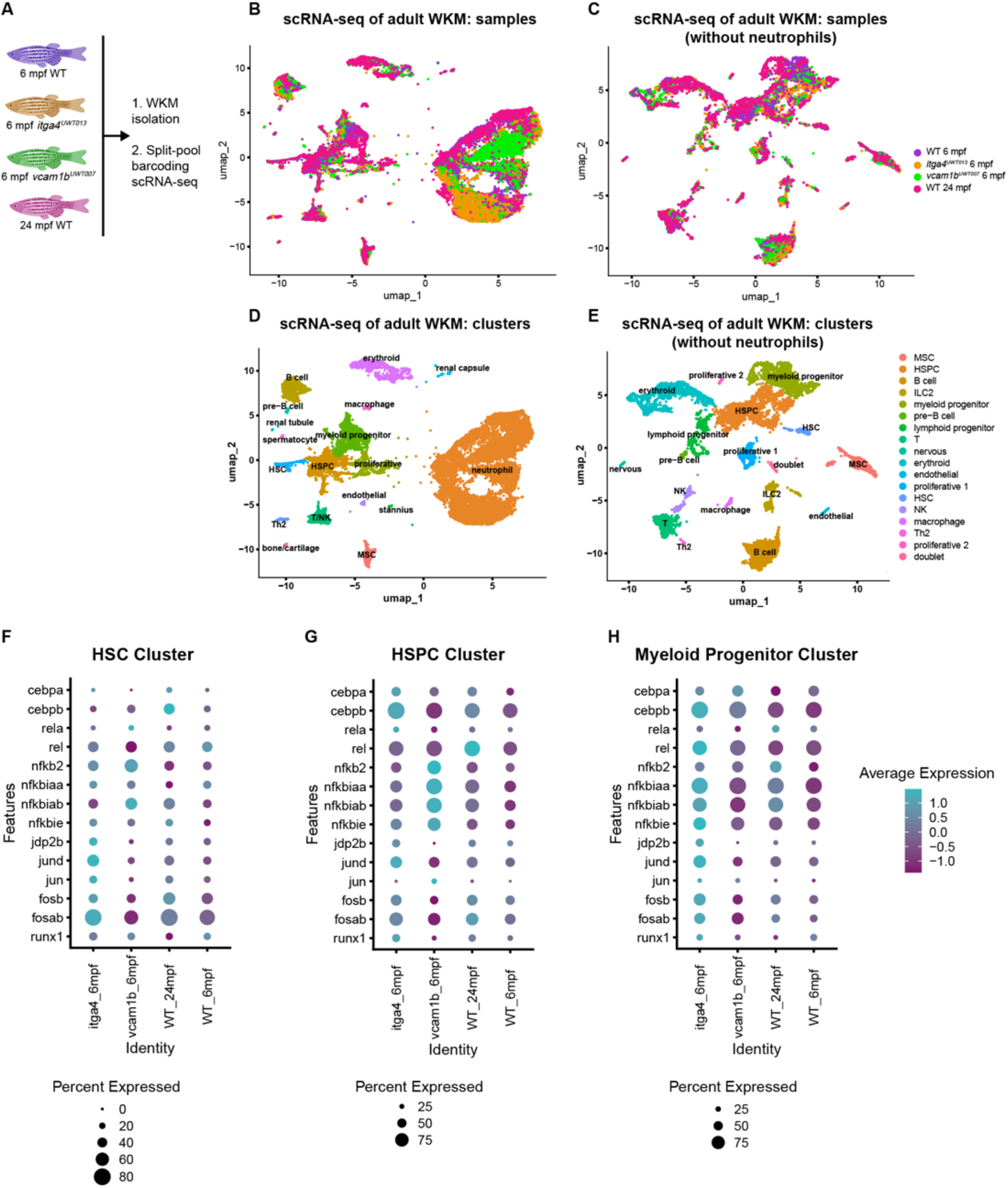
Adult WKM scRNA-seq analysis shows *itga4* mutant HSCs maintain an inflammatory profile. **A)** The workflow for split-pool barcode scRNA-seq of adult WKM, excluding erythrocytes via fluorescence-activated cell sorting (FACS) or red blood cell (RBC) lysis (created with Biorender). One biological replicate is WKM from one adult fish. Biological replicates for each condition: n=4 each for 6 mpf WT, 6 mpf *itga4^UWT013^*, and 24 mpf WT; n=3 for 6 mpf *vcam1b^UWT007^*. Sample processing, library prep, and sequencing occurred over three independent experiment days. All replicates were included for analysis. Total cells from all replicates: 6 mpf WT n=5,202; 6 mpf *itga4^UWT013^* n=8,149; 24 mpf WT n=7,352; 6 mpf *vcam1b^UWT007^* n=5,033. **B-C)** Color-coded UMAP projection for all samples after batch correction (key on right): purple=6 mpf WT; orange=6 mpf *itga4^UWT013^*; green=6 mpf *vcam1b^UWT007^*; pink=24 mpf WT, **B)** with, or **C)** without neutrophils. **D-E)** UMAP projection for all clusters after batch correction, with color coding based on cluster identity (key on right), **D)** with, or **E)** without neutrophils. **F-H)** Bubble plots showing transcript expression of AP-1 and NFκB pathway factors in the HSC, HSPC, and Myeloid Progenitor clusters for each condition. Expression levels of *cebpa*, *cebpb*, and *runx1* were also included for each cluster. Each plot has corresponding legends below. Dot color indicates expression level (light blue=high expression, purple=low expression). Dot size indicates percent of cells in each cluster that express the marker and average expression is log2 normalized. **F)** HSC Cluster; **G)** HSPC Cluster; **H)** Myeloid Progenitor Cluster. See also **Figure S7 and Table S4**

We used split-pool barcoding scRNA-seq as an unbiased approach to capture both hematopoietic and niche cells. We excluded red blood cells and collected all other WKM cell types in the expected proportions^90, 91^, with the largest proportion being neutrophils (Fig. 5B,D; S7A). We observed batch effects within the neutrophil cluster so removed them before further analysis (Fig. 5C,E). We performed pseudo-bulk analysis on HSC, HSPC, and myeloid progenitor clusters separately in each of the different backgrounds (i.e., young WT, *itga4* and *vcam1b* mutants, and aged WT; Fig. 5F-H; Table S4). Based on our analysis of developmental *itga4* mutant HSPCs, we looked specifically at AP-1 and NF-κB pathway factors, and included markers of myelopoiesis (*cebpa, cebpb*) and HSPCs (*runx1*). The proportion of cells that expressed these markers was largely unchanged across ages and backgrounds. However, young *itga4* mutant adult HSCs, HSPCs, and myeloid progenitors had higher expression of AP-1 factors (Fig. 5F-H), as we observed during development (Fig. 1,2). In HSPCs, it was exclusively young *vcam1b* mutant adults that had increased expression of NF-κB pathway factors (Fig. 5G), suggesting the *itga4* and *vcam1b* mutant phenotypes may not be equivalent. Aged HSCs had higher expression of *cebpb* that is consistent with the myeloid-biased output of aged hematopoiesis^92^.

We performed a module score analyses to compare adult HSPCs from each condition against publicly available Human Hallmark Gene Sets from the MolSigDB database^93^ for the following pathways: Interferon Alpha Response, Interferon Gamma Response, and Myc Targets V1 (Fig. S7B-D). A greater proportion of 6 mpf *itga4* mutant HSPCs scored positively against the Interferon Alpha and Gamma Response module sets compared to all other conditions (Fig. S7B,C). A h<u>igher proportion</u> of 6 mpf *itga4* mutant and *vcam1b* mutant HSPCs scored positively for the Myc Targets V1 module set compared to young and old adult WT HSPCs (Fig. S7D). Overall, scRNA-seq analysis found similarities between larval and adult *itga4* mutant HSPCs, with both exhibiting upregulation of inflammatory and AP-1 pathways. However, we do not know if this phenotype persists from development into adulthood, or if it arises separately in the adult marrow from reduced Itga4-Vcam1 adhesion and Itga4-dependent signaling.

### Young adult *itga4* mutant HSPCs have a distinct chromatin landscape that is enriched for GATA TF binding motifs

To determine if *itga4* mutant HSPCs have an altered chromatin landscape in adulthood, we used bulk ATAC-seq to compare sorted Runx:mCherry+ HSPCs from young 6 mpf WT, *itga4* mutant, and *vcam1b* mutant, as well as aged 24 mpf WT adult WKM (Fig. 6A). We found there was “gain” and “loss” of accessibility to regions of chromatin in young *itga4* mutant HSPCs relative to young WT HSPCs (Fig. 6B). The genome-wide distribution of chromatin accessibility for promoters, coding sequences, intergenic regions, etc. was similar in HSPCs for all genetic backgrounds and ages (Fig. S8A). Pairwise comparisons of chromatin accessibility peaks were performed between young WT adult and each background (i.e., young adult *itga4* and *vcam1b* mutant, and aged WT adult) (Fig. S8B-D; Table S5). Venn diagrams show young adult *itga4* HSPCs have the most unique peaks of each background, when compared to young WT adult (Fig. S8C; Table S5). While *itga4* and *vcam1b* mutant HSPCs had some common peaks relative to young WT adult (Fig. S8C,D), most peaks were unique, suggesting disruption of either side of the Itga4-Vcam1 axis was not equivalent, and HSPC phenotypes in each background are distinct.

**Figure 6.**
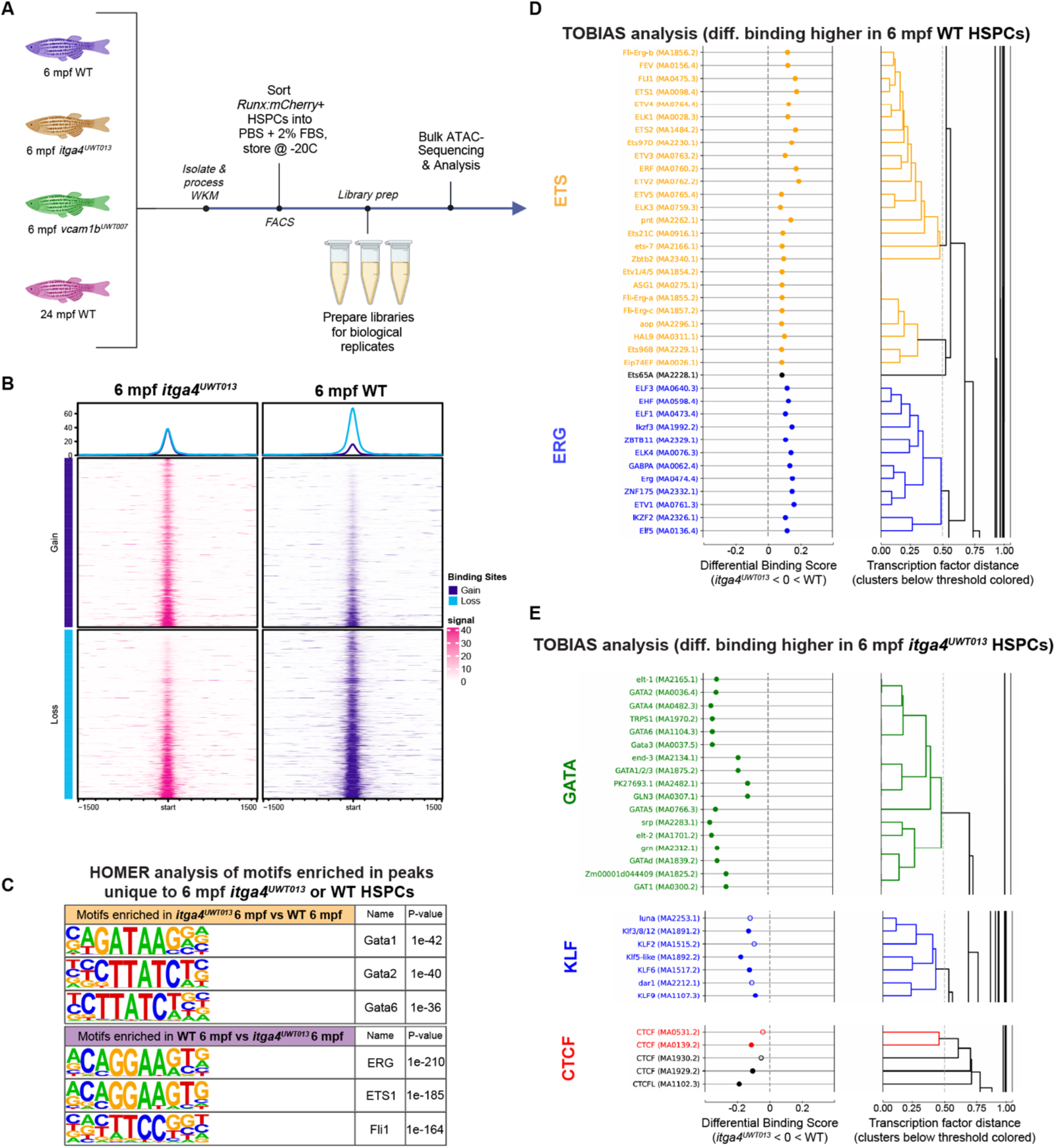
Bulk ATAC-seq of adult HSPCs reveals an *itga4* mutant chromatin landscape enriched for GATA TF binding motifs. **A)** Bulk ATAC-seq workflow for Runx:mCherry+ HSPCs sorted from adult WKM, excluding erythrocytes via RBC lysis (created with Biorender). One biological replicate is WKM from one adult fish, with n=3 biological replicates per condition. Sample processing, library prep, and sequencing occurred during one experiment. All replicates were included for analysis. Total cells from all replicates: 6 mpf WT n=58,099; 6 mpf *itga4^UWT013^* n=62,480; 6 mpf *vcam1b^UWT007^* n=35,438; 24 mpf WT n=61,821. **B)** Metaplots (top) and heatmaps (bottom) showing differential chromatin accessibility between *itga4^UWT013^* and WT HSPCs (plots centered on TSS). “Gain” and “Loss” means increased or decreased accessibility, respectively, in regions of chromatin from 6 mpf *itga4^UWT013^* compared to 6 mpf WT. Upper: Metaplot average accessibility signal (purple “gain” curves and blue “loss” curves). Lower: Heatmap accessibility signal from all peaks (stronger color saturation=greater accessibility; left/pink=*itga4^UWT013^*; right/purple=WT). **C)** Top predicted TF binding motifs from HOMER known motif analysis of uniquely accessible peaks in each condition. Representative top 3 motifs and p-values are shown for the GATA family enriched in 6 mpf *itga4^UWT013^* (upper), and the ETS family enriched in 6 mpf WT (lower). **D-E)** Results from TOBIAS TF binding prediction analysis comparing 6 mpf WT and 6 mpf *itga4^UWT013^*HSPCs. Hierarchical clustering based on motif similarity (color=similar; black=unique). **D)** TFs predicted to gain binding sites in 6 mpf WT compared to 6 mpf *itga4^UWT013^*, grouped into motif families representing ETS and ERG factors. **E)** TFs predicted to gain binding sites in 6 mpf *itga4^UWT013^*compared to 6 mpf WT, grouped into motif families representing CTCF, KLF, and GATA factors. See also **Figure S8 and Table S5**.

We performed HOMER motif enrichment analysis on uniquely accessible regions from each pairwise comparison (Fig. 6C; Table S5). Young adult WT HSPCs feature predominant enrichment of ERG/ETS1 binding motifs when compared to young adult *itga4* mutant HSPCs (Fig. 6C; Table S5). This was consistent with predicted TF footprinting using TOBIAS that also showed enriched ERG/ETS1 motifs (Fig. 6D). The ETS factor *Erg* has been shown to increase in expression level from fetal to adult stages and is required for HSPC maturation and implementation of adult regulatory programs^94^. Accessible regions in young adult *itga4* mutant HSPCs compared to young WT were enriched for CTCF, KLF, and GATA motifs using both HOMER and TOBIAS analysis (Fig. 6C,E; Table S5). These data suggest the Itga4-Vcam1 axis is also involved in maintaining chromatin accessibility in adult HSPCs, however, there may be shifting requirements for TF regulatory programs between 5 dpf *itga4* mutant and WT larval HSPCs, and young adult *itga4* mutant and WT WKM HSPCs. The persistent dysregulation of the chromatin landscape and TF regulatory networks may be a continuation of developmental changes in *itga4* mutant HSPCs, or from diminished attachment to the adult niche.

### Young adult *itga4* mutant HSPCs are inflammatory, proliferative, and myeloid-biased

To understand the phenotypic consequences of altered gene regulatory programs in adult *itga4* loss of function HSPCs, we wanted to consider if there was a lineage bias in HSPCs from young *itga4* mutant WKM compared to WT. After red cell lysis, we analyzed Runx:mCherry+ HSPCs based on forward and side scatter that can distinguish between lymphoid, precursor, and myeloid cells in WKM^95^ (Fig. 7A). We found the majority of Runx:mCherry+ HSPCs in the lymphoid gate from young adult WT WKM based on their size and granularity; this was expected as we showed it previously^30^. However, in young adult *itga4* mutant WKM compared to young adult WT, there was an increase in the proportion of Runx:mCherry+ HSPCs found in the myeloid progenitor and precursor gates, at the expense of HSPCs in the lymphoid progenitor gate (Fig. 7A-D).

**Figure 7.**
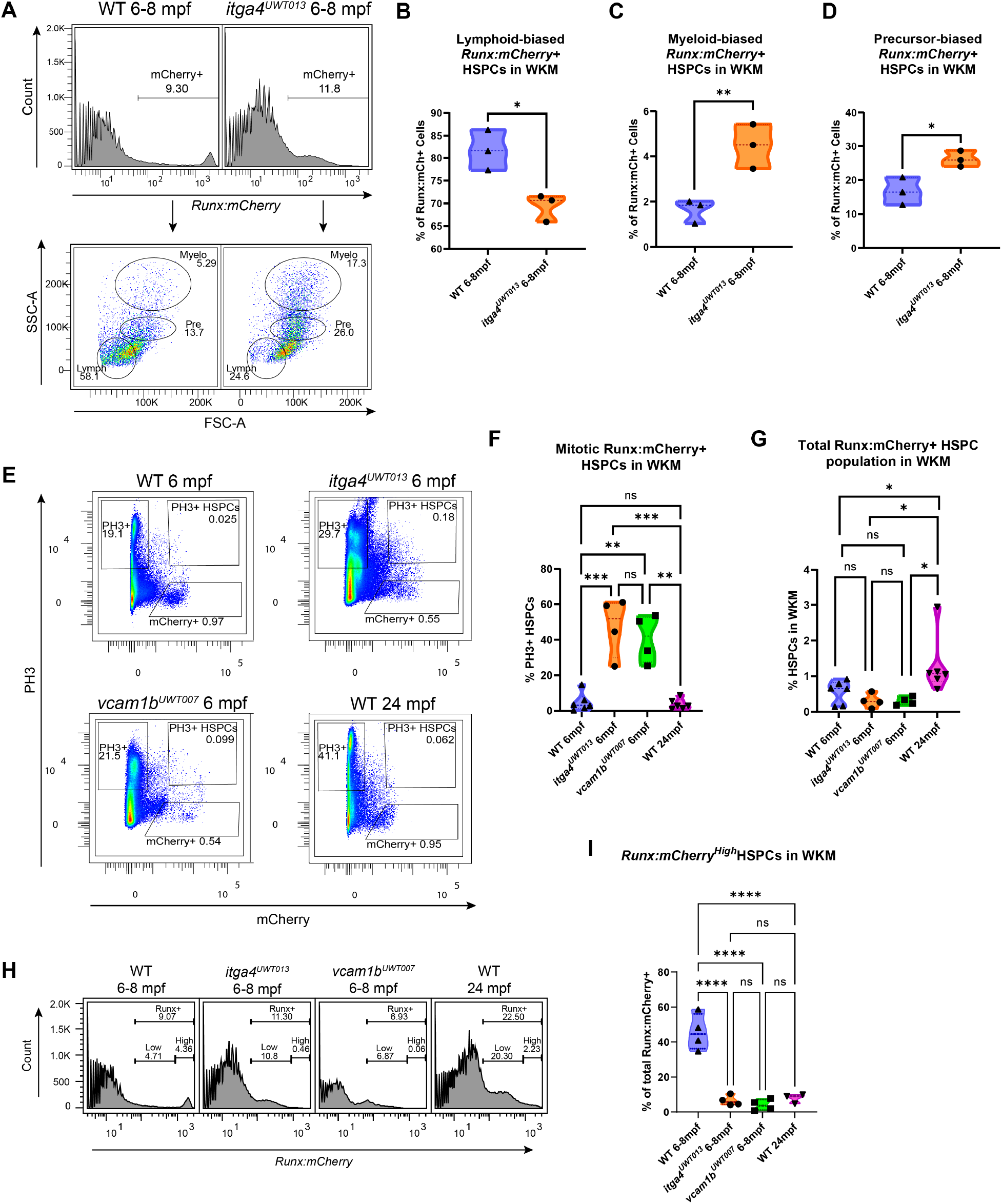
Adult WKM flow cytometry analysis shows young adult *itga4* mutant HSPCs are inflammatory, proliferative, and myeloid-biased. **A-D**) Runx:mCherry+ HSPC lineage bias from 6-8 mpf WKM by forward and side scatter. Replicates: n=3 for both conditions. **A)** Representative gating strategy. HSPC lineage biases: **B)** Lymphoid: WT median=81.5 (IQR=8.98); *itga4^UWT013^*median=70.7 (IQR=5.60); *p=0.0122. **C)** Myeloid: WT median=1.85 (IQR=0.973); *itga4^UWT013^* median=4.51 (IQR= 1.98); **p=0.00748; **D)** Precursor: WT median=16.5 (IQR=8.15); *itga4^UWT013^*median= 25.9 (IQR=4.7); *p=0.0179. **E-G)** The proportion of dividing PH3+ HSPCs by flow cytometry. Replicates: n=6 for 6 mpf WT and 24 mpf WT; n=4 for 6 mpf *itga4^UWT013^* and 6 mpf *vcam1b^UWT007^*. **E)** Representative gating strategy. **F)** Mitotic HSPCs: 6 mpf WT median=3.16 (IQR=6.94); 6 mpf *itga4^UWT013^* median=51.8 (IQR=30.7); 6 mpf *vcam1b^UWT007^* median= 42.1 (IQR=25.3); 24 mpf WT median= 2.81 (IQR=5.10). Significant comparisons: 6 mpf WT vs 6 mpf *itga4^UWT013^* (***p=0.0005); 6 mpf WT vs 6 mpf *vcam1b^UWT007^* (**p=0.0088); 6 mpf *itga4^UWT013^*vs 24 mpf WT (***p=0.0002). **G)** Percent Runx:mCherry*+* HSPCs from live cells. HSPC percentage: 6 mpf WT median=0.652 (IQR=0.661); 6 mpf *itga4^UWT013^*median= 0.286 (IQR=0.369); 6 mpf *vcam1b^UWT007^* median=0.286 (IQR=0.218); 24 mpf WT median=1.047 (IQR=0.326). Significant comparisons: 6 mpf *itga4^UWT013^* vs 24 mpf WT (*p=0.0105); 6 mpf *vcam1b^UWT007^* vs 24 mpf WT (*p=0.0214); 6 mpf WT vs 24 mpf WT (*p=0.0356 with extreme observation included in 24 mpf WT group; extreme excluded: p=0.0584). **H-I)** Proportion of Runx:mCherry^High^ HSPCs in WKM. Replicates: n=4 for all conditions, except 24 mpf WT (n=3). **H)** Representative gating of Runx:mCherry+ HSPCs. **I)** Runx:mCherry^High^ percentage from total HSPCs: 6 mpf WT median=44.5 (IQR=19.8); 6 mpf *itga4^UWT013^* median=5.70 (IQR=5.29); 6 mpf *vcam1b^UWT007^* median=3.65 (IQR=6.02); 24 mpf WT median=8.97 (IQR=5.10). Significant comparisons: 6 mpf WT vs 6 mpf *itga4^UWT013^* (****p<0.0001); 6 mpf WT vs 6 mpf *vcam1b^UWT007^* (****p<0.0001); 6 mpf *itga4^UWT013^* vs 24 mpf WT (****p<0.0001). Parent gates: Live single cells. Biological replicates: WKM from one fish; RBC lysis; three experiment days. Statistical analyses: all replicates included; percentages analyzed using generalized linear models for over-dispersed binomial data; p values for comparisons among groups adjusted for multiplicity using single-step procedure that relies on multivariate t-distribution for all pairwise comparisons^139^. See also **Figure S9**.

Next, we assessed *in vivo* cell proliferation and lineage output in the WKM of young 6 mpf WT, *itga4* and *vcam1b* mutants, and 24 mpf WT (Fig. 7E-G). There was a higher proportion of mitotic phosphorylated histone H3-positive HSPCs in 6 mpf *itga4* and *vcam1b* mutants compared to both 6 and 24 mpf WT (PH3+;Runx:mCherry+; Fig. 7E,F, S9A). In aged 24 mpf WT WKM, it was the non-HSPC fraction was more mitotic (PH3+;Runx:mCherry-; Fig. S9B). Interestingly, only aged 24 mpf WT adults had a higher overall proportion of HSPCs in WKM, even though that population was not more mitotic (Fig. 7F,G). This is consistent with one of the hallmarks of aged hematopoiesis in mammals that shows a higher frequency of HSPCs^92^.

To consider putative HSCs versus the broader HSPC population that includes more differentiated progenitors, we focused on Runx:mCherry^high^ cells that our scRNA-seq data found clustered together and annotated as HSCs (Fig. S2D). This is consistent with our previous observation that Runx:mCherry^high^ cells are more stem-like and give rise to Runx:mCherry^low^ progenitors^30^. In young adult WKM we found Runx:mCherry^high^ HSCs were decreased and Runx:mCherry^low^ progenitors were increased in *itga4* and *vcam1b* mutants compared to WT (Fig. 7H,I). It is possible the higher mitotic rate in mutant HSPCs (Fig. 7F), may be leading to HSC exhaustion. Interestingly, aged WT WKM also exhibited a similar decrease in Runx:mCherry^high^ HSCs and increase in Runx:mCherry^low^ progenitors (Fig. 7H,I), even though aged Runx:mCherry+ HSPCs do not have a higher mitotic rate (Fig. 7F). Another hallmark of aged hematopoiesis is an increase in myeloid-biased HSCs^92^ that may explain the increase in Runx:mCherry^low^ progenitors (Fig. 7H). Together, our results show that HSPCs in young adult *itga4* mutants are myeloid biased, more mitotic, and have fewer HSC-like Runx:mCherry^high^ cells.

## Discussion

The programming of HSPCs as they transition from DA emergence, to CHT or FL lodgement, and finally population of the marrow, follows changing metabolic, proliferative, and differentiation requirements. After HSCs transition through multiple states and locations they ultimately shift from a proliferative state to quiescence^3, 6, 7, 9, 22, 25, 96^. Recent work has shown that HSPCs in the CHT or FL may not be as proliferative as previously thought, and instead undergo maturation and differentiation^48, 97, 98^. Here we find that *itga4* mutant HSPCs after CHT stage at 5 dpf have activated inflammatory pathways and retain an immature phenotype like HSPCs that have just emerged from the DA^33, 34, 36, 37, 83, 84^. This suggests that contact with the CHT niche, and possibly Itga4 signaling itself, is necessary for HSPC maturation by downregulation of inflammatory pathways, chromatin remodeling, changing TF regulatory programs, and ultimately transition to quiescence (Fig. 8).

**Figure 8.**
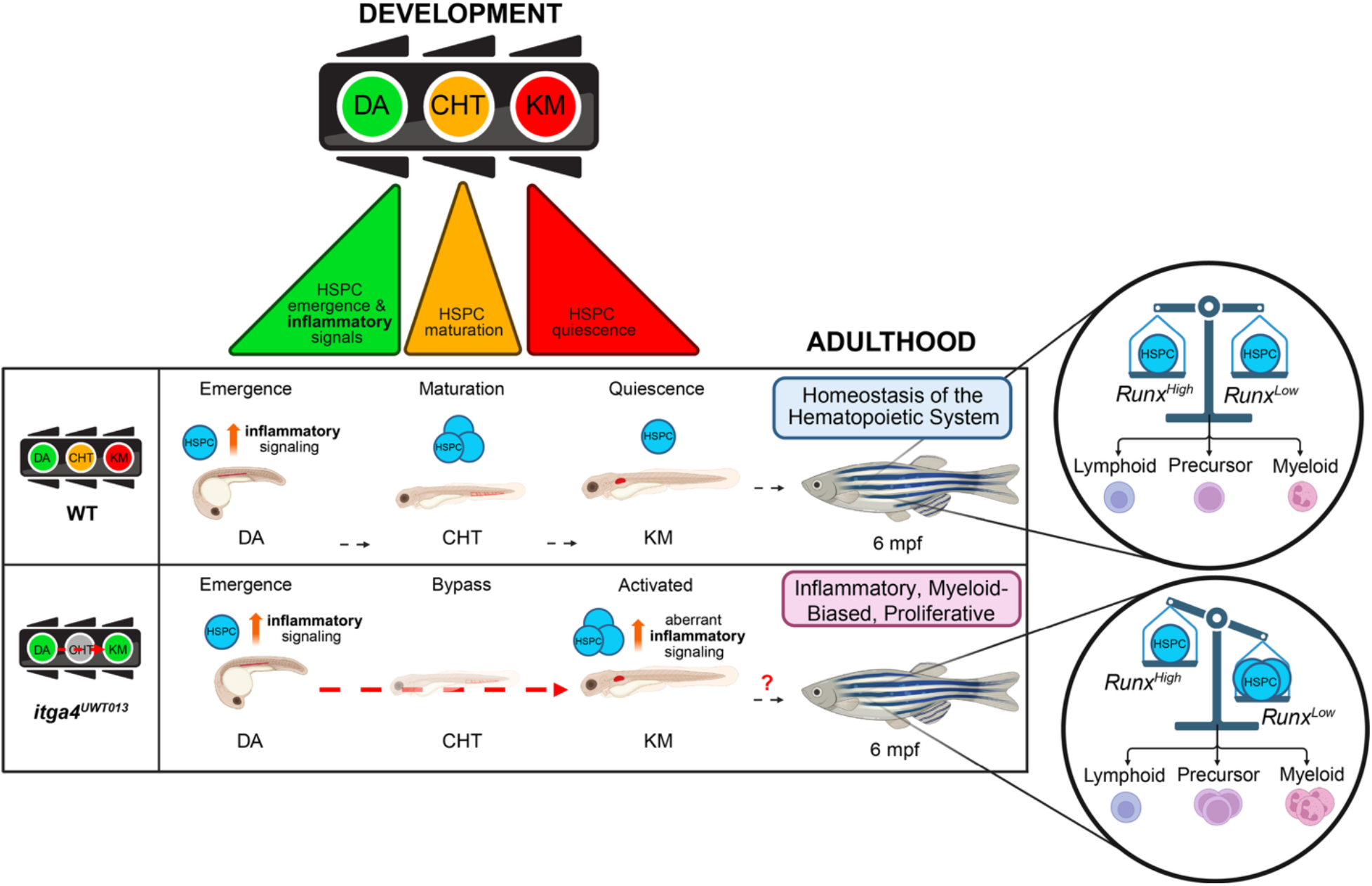
Model of HSPC maturation in WT and itga4 mutant embryos. A proposed “traffic light” model for how the developmental CHT niche promotes HSPC maturation (created with Biorender). From left to right: The “green light” stage features pro-inflammatory signals and HSPC emergence; the “yellow light” stage indicates down regulation of pro-inflammatory signals and the start of HSPC maturation; the “red light” stage marks the transition to HSPC quiescence. In the *itga4* mutant model, bypassing the CHT niche maintains HSPCs in an active “green light” state through development. Young adult *itga4* mutant HSPCs are more proliferative and share some features with aged HSPCs, such as high levels of inflammatory pathway genes and a myeloid lineage bias.

Chronic low-level inflammation, including aging-associated “inflammaging”, has a significant impact on hematopoiesis and HSC function^89, 97, 99^. In our study we included aged (24 mpf) zebrafish to compare their hematopoietic phenotypes to young adult *itga4* mutants that share characteristics with the aging phenotype, such as a myeloid bias and an inflammatory profile^100,101^. One distinct difference we found is that young adult *itga4* mutant HSPCs have elevated NF-κB activity, but are mitotically active, compared to young and aged WT WKM that only have a small percentage of dividing HSPCs. Although NF-κB activity is higher in aged HSCs^102^, the intrinsic and extrinsic effects of NF-κB signaling may differ by promoting quiescence and myeloid bias, respectively^76^. Likewise, in *itga4* mutant zebrafish HSPCs, it may be that either proliferation or myeloid bias, but not both, are coupled to inflammatory pathway activity.

During development, changing levels of NF-κB activity are linked to proliferation as HSPCs go through EHT in the DA^35^. In IκBα (*Nfkbia* inhibitor of NF-κB) mutant mouse embryos there is sustained NF-κB activity that prevents HSC maturation in the FL, and as a result HSCs retain an emergent DA phenotype^41^. This elevated NF-κB activity in HSCs promotes quiescence and diminishes stem cell function^41, 76^. Mechanistically, NF-κB regulates RA pathway targets that are associated with HSC dormancy^41, 80^. Consistent with maturation and the transition to quiescence during development, we observed establishment of a RA regulatory program by motif analysis of chromatin accessible regions in WT 5 dpf HSPCs (Fig. 3). However, in *itga4* mutant 5 dpf HSPCs we found enrichment of motifs for ETS and AP-1 factors. Interestingly, these factors are associated with emergent, newly formed HSPCs in the DA. The zebrafish homolog of human ETS factor SPI1, *spi2*, is required for lymphoid/myeloid-primed HSPCs^103^. AP-1 factors are particularly important for early and emergent HSPCs^83, 84^. These results suggest that Itga4 function and CHT contact are required for HSPC transitions through development to alter their chromatin landscape and change gene regulatory programs.

During EHT there is a switch from glycolysis to oxidative phosphorylation^104, 105^. In zebrafish, energy metabolism is dynamic during EHT and dependent on glucose and oxygen availability^39, 106^. ROS is high in cycling HSCs^107^ and FL HSCs typically have higher oxidative phosphorylation (OXPHOS) genes to support their metabolic demands^15^. However, in FL HSCs mitochondrial function is uncoupled from proliferation, and instead is required for differentiation^108^. Elevated NF-κB activity in FL HSCs not only delays maturation but also has the effect of down-regulating OXPHOS and MYC-target pathways^41^. These characteristics are strikingly similar to *itga4* mutant HSPCs at 5 dpf analyzed by bulk RNA-seq that also have down-regulated OXPHOS and MYC-target pathways (Fig. 1), and *itga4* mutant phenotypic HSCs at 5 dpf analyzed by scRNA-seq that have a more immature profile (Fig. 2). Together, this shows Itga4 function and CHT contact not only changes gene regulatory programs of HSPCs, but also their metabolic profile as they mature through proliferative and ultimately quiescent states.

There are several outstanding questions that will need to be addressed by future studies. First, we do not know if the phenotype of *itga4* mutant HSPCs in adulthood results from inflammatory signaling that is continuously elevated from early development, or if it results from disrupted ITGA4-VCAM1 signaling and adhesion in the adult WKM. In mice, conditional deletion of *Itga4* in adult HSCs impairs homing to the BM niche and has some impact on stem cell engraftment^53, 109^. The role of Itga4 in adhesion to the BM niche is further demonstrated by administration of the small molecule inhibitor of VLA-4, BIO5192, that mobilizes HSPCs^110^. Therefore, it is possible that the adult zebrafish *itga4* mutant HSPC phenotype could result from destabilized adhesion to the niche.

We observed that young adult *itga4* mutant HSPCs share some of the hallmarks of aged HSPCs, including a myeloid bias^101^. Importantly, it will need to be determined if *itga4* mutants have clonal hematopoiesis that is either lost during development or in adulthood. It will be necessary to uncouple the function of Itga4 in adhesion of HSPCs to the niche, or its function in “outside-in” signaling intrinsically in HSPCs. It was previously shown that another integrin, Itgav, has a signaling role in HSCs^111^. Chromatin level and gene expression changes in both developmental and adult *itga4* mutant HSPCs indicate intrinsic programming changes that result from impaired niche contact. Although *itga4* mutant HSPCs become more proliferative, they also have elevated inflammatory pathways and altered metabolic function. Therefore, reversal of adult quiescence to favor expansion for translational purposes should be carefully considered. Maintaining HSPCs in an immature state could have unintended consequences and ultimately create a stem cell pool with diminished function and increased susceptibility to pathologies.

## Methods

### Ethics declarations and approval for animal experiments

Zebrafish (*Danio rerio*) were maintained at the University of Wisconsin-Madison or the Iowa State University in accordance with their respective Institutional Animal Care and Use Committee guidelines. All zebrafish experiments were performed according to Iowa State University Animal Care and Use Committee IACUC-20-025 and IACUC-20-024 approved protocols, and in compliance with ARRIVE guidelines^112^, and the American Veterinary Medical Association (2020) and NIH guidelines for the humane use of animals in research.

### Zebrafish Husbandry

Zebrafish were maintained in a circulating aquarium system at 28°C with a 14/10 hour light/dark cycle, and handled as described in standard protocols^113^. Embryos were raised in 1x E3 medium (5 mM NaCl, 0.17 mM KCl, 0.33 mM CaCl_2_, 0.33 mM MgSO_4_) with methylene blue and maintained at 28°C. Sex cannot be determined in zebrafish at larval stages^114^, and therefore was not considered as a biological variable for experiments at 5 dpf timepoints. In adults (6-24 mpf), we analyzed an equivalent number of males and females for each condition when possible (indicated in figure legends). No significant differences were found between adult males and females in this study.

### Zebrafish Lines

Zebrafish lines used in this study were: *Tg(Runx:mCherry)*^30^; *Tg(EF1α:mAG-zGem(1/100))^rw0410h^*, referred to as *zFUCCI*^85^; *Tg(6xHsa.NFKB:d2mRFP1)^is503^*, referred to as *NFKB:d2mRFP*^35^; *Tg(−6.0itga2b:eGFP)^la2^*, referred to as *cd41:eGFP*^86^. The *itga4^UWT013^* and *vcam1b^UWT007^* mutant alleles were established at the University of Wisconsin-Madison as previously described ^62^.

### Genotyping Zebrafish Lines

The *itga4^UWT013^* and *vcam1b^UWT007^* lines were genotyped using the primers below (5’-3’ orientation). PCR products from *itga4* mutants were digested with MscI, and *vcam1b* mutants with DraI to distinguish WT, heterozygous, and homozygous siblings:

#### itga4^UWT013^

FWD: ACACTCTTTCCCTACACGACGCTCTTCCGATCTGGCTGAAATGCTCCATTCAT

REV: GTGACTGGAGTTCAGACGTGTGCTCTTCCGATCTGATGGCTCCAGGACTTTTGA

#### vcam1b^UWT007^

FWD: ACACTCTTTCCCTACACGACGCTCTTCCGATCTCGGTCCAGACAAACTTCACC

REV: GTGACTGGAGTTCAGACGTGTGCTCTTCCGATCTTGCAACTGAACAAGCATGAA

### Morpholino injections

*itga4* MO (5′-CTCCAGTGAGTTGTGATGAAATCAT-3′) was diluted with nuclease free water to a concentration of 100 µM. 1% concentration of phenol red solution (MilliporeSigma; #P0290) was used as a pH indicator. A final 500pL volume was injected into each embryo at the 1-2 cell stage.

### Head/Trunk and Tail Isolations

*Tg(NFKB:d2mRFP; cd41:eGFP)* embryos were dechorionated with pronase (Millipore Sigma, #11459643001) and anesthetized in 1% tricaine (Pentair, #TRS5). The tail region of each embryo was surgically isolated (posterior to the urogenital canal) using scalpel blades (Roboz Surgical Store, #RS-9801-15).

### Tissue Dissociation and Single Cell Suspension

Larvae (n≥100) were pooled for each sample at 5 dpf to be dissociated into a single-cell suspension, based off previously published protocol^115^. Mechanical homogenization (step 2.2 of the published protocol) was performed with disposable pellet pestles (Fisherbrand, #121-141-364) in alternation with harsh pipetting. Samples underwent stages of “rest” on a heating block set to 35°C during the time needed to complete each round of mechanical homogenization for all samples. Additions to the protocol include extra 40 μm filtrations (VWR, #76327-098) at the end of the dissociation stage and after resuspending the pellet in the washing stage (steps 2.3 and 3.1 of the published protocol, respectively). This adjusts the protocol to have three total filtrations and allows for filtration before each centrifugation step. Centrifugation settings were also adjusted to 10 minutes at 300g at room temperature. The final filtration and wash step (step 3.2 of published protocol) was performed with 1x PBS supplemented with 2% FBS and completed in 5mL tubes for flow cytometry applications (Corning, #352054).

*Tg(NFKB:d2mRFP; cd41:eGFP)* embryos were screened for fluorescence and dissociated as previously described (PMID:<u>35704898</u>). Briefly, double transgenic embryos were dechorionated with pronase (MilliporeSigma, #11459643001) and treated with embryo water containing 0.003% 1-Phenyl-2-thiourea (MilliporeSigma, #P7629) from 22hpf to inhibit production of melanin. At the target developmental stage, embryos were dissociated with 50µg/mL Liberase TM (MilliporeSigma, #5401127001) in Dulbecco′s Phosphate Buffered Saline solution with Ca^2+^ and Mg^2+^ (MilliporeSigma, #D8662), assisted by gentle vortex at 28.5 °C for 20-30 minutes. The resulting cell suspension was filtered through a 30 µm nylon mesh (Fisher Scientific, #NC9084441) using a 21-gauge syringe (Fisher Scientific, #BD 309624) and stained with SYTOX™ Red (Life Technologies, #S34859) to exclude dead cells.

Dissection and single cell suspension of adult WKM followed previously published methods^116^.

### Flow Cytometry and Cell Sorting

Upon obtaining single cell suspension, cells were counted with a haemocytometer and dead cells were stained with trypan blue at a dilution of 1:10. Cells were suspended in 1x PBS supplemented with 2% FBS and analyzed via flow cytometry using a BD LSR Fortessa or sorted via fluorescent activated cell sorting (FACS) with BD FACS Attune or Aria II (BD Biosciences). Dead cells were excluded using Dapi. For DNA sequencing applications, cells were sorted via FACS into 1x PBS supplemented with 2% FBS. For RNA sequencing applications, cells were sorted via FACS into Trizol LS (Invitrogen, #10296010). Data was analyzed with FlowJo software (version 10.3 or 10.10). The cell suspension resulting from *Tg(NFKB:d2mRFP; cd41:eGFP)* embryos was analyzed by flow cytometry in a BD FACSMelody Cell Sorter (BD Biosciences) with FACSChorus 2.0, version 1.1.20.0 and FlowJo software (version 10.3 or 10.8.1, Tree Star).

### Bulk RNA-seq and data analysis

Runx:mCherry+ HSPCs were sorted via FACS directly into Trizol LS (Thermo Fisher Scientific, #10296028) and stored at -80 °C until all replicates were collected. Total cells from all replicates: n=46,712 WT and n=6,413 *itga4^UWT013^*. Once all replicates were collected, samples were brought to room temperature and the protocol for total RNA isolation was completed based on Invitrogen protocol Cat. No. 10296-010. Isolated RNA was sent to the Biotechnology Center at the University of Wisconsin-Madison for quality control analysis via TapeStation, library preparation, and sequencing. RNA libraries for RNA-seq were prepared using Takara SMART-Seq v4 Ultra Low Input RNA/NexteraXT (polyA enrichment) following manufacturer’s protocols.

Reads underwent adapter trimming with “skewer” v.0.2.2b^117^, then were mapped to a custom GRCz11 reference genome with “STAR” v.2.5.3a^118^. The GRCz11 reference genome was customized to include inserted transgenic reporter constructs (e.g. *Runx:mCherry*). Mapped reads were quantified with “RSEM” v.1.2.31^119^ and processed via “edgeR” v.3.34.0^119^ to independently filter samples with the *filterByExpr* function to remove genes with less than 10 read counts, normalize each sample by the method of trimmed mean of M-values^120^, and perform analysis of differentially expressed genes between samples while controlling the false discovery rate^121^. Finally, transcripts were annotated with the “Ensembl” annotation system^122^ and DEGs underwent downstream analysis with GSEA^123^.

### Bulk ATAC-seq and data analysis

Runx:mCherry+ HSPCs were sorted via FACS directly into PBS supplemented with 2% FBS for downstream processing. Total cells from all replicates: n=23,524 WT and n=22,526 *itga4^UWT013^*. Libraries for ATAC-seq were prepared based off previously published protocol for ATAC-seq^124^. Primer sequences used to generate the libraries were referenced from previously published protocol^125^, with additional size selection of fragments and library cleanup using AmPure XP Beads (Beckman Coulter, #A63880). The number of PCR cycles for library amplification was optimized for organism and cell type, which deviated from the referenced published protocols. Unpooled libraries were sent to the Biotechnology Center at the University of Wisconsin-Madison for quality control analysis via TapeStation and sequencing.

Sequences were analyzed by ENCODE ATAC-seq pipeline (v2.2.3) with customized GRCz11 genome with additional DsRed2, ZsGreen, mCherry, and EGFP gene sequences. The conservative set selected by IDR (Irreproducible Discovery Rate) were used for Downstream analysis. DiffBind^126^ (v3.16.0; DOI:<u>10.18129</u>) was used to call differential accessible sites (DAS). ATAC-Seq peaks were annotated nearest gene symbols by ChIPpeakAnno v3.40.0^127^. The motif enrichment analysis was performed by TFEA v1.1.2^81^. The motifs enrichment was also performed for the DAS filtered with FoldChange>1.1 and P.value<0.05 by Homer v5.1^79^. The promoter and gene body usage analysis were performed by LIMMA v3.62.2^128^ voom for the reads counts in each region. The promoters were defined as upstream 1K to downstream 500bp from the TSS. The gene body was defined as gene ranges without the promoter.

TF footprinting analysis was performed using TOBIAS v0.17.3^129^. To generate a common reference set of accessible chromatin regions for footprinting analysis, the IDR optimal peak sets from the wild-type and mutant samples were combined, sorted, and merged using bedtools v2.31.1^130^ merge to create a consensus peak set. Peak annotations and genomic regions of interest were generated using the TOBIAS region preparation utilities with the parameters -- feature CDS, --feature_anchor start, and --distance 20000 10000, defining regions spanning 20 kb upstream and 10 kb downstream of coding sequence start sites. TF binding motifs were obtained from the JASPAR2026 CORE^131^ non-redundant position frequency matrix collection. TOBIAS ATACorrect was used to correct Tn5 transposase sequence bias for each sample, followed by FootprintScores to calculate genome-wide footprint scores. Differential TF binding between wild-type and mutant samples was assessed using BINDetect, which integrates corrected footprint scores with motif occurrences to identify TFs with differential predicted binding occupancy between the two conditions.

### scRNA-seq and data analysis

Runx:mCherry+ HSPCs were sorted via FACS directly into 0.2mL PCR strip tubes with PBS supplemented with 2% FBS for a “sort and load” protocol directly into the library preparation stage. Samples were then immediately loaded onto a 10X Genomics 3’ v3.1 chip for processing according to manufacturer’s instructions. Libraries for scRNA-seq were prepared at the Biotechnology Center at the University of Wisconsin-Madison utilizing reagents and protocols supplied with the 10X Genomics 3’ v3.1 kit under standard conditions.

The “cell ranger” v.9.0.0 pipeline (10X Genomics) was utilized for initial processing and alignment to the custom GRCz11 reference genome. Further data processing was completed via R v.4.5.1, where cells with low detected features or high percent of mitochondrial reads were removed by a threshold of 2 median absolute deviations from the median and the package “SoupX” was used to remove ambient RNA^132^. Data was normalized using size factor/log transformation and quality control was performed to remove low-quality cells and doublets. The package “batchelor” was used to reduce batch effect on reduced dimensions^133^. Clustering was performed with three levels of resolution before assigning cell types via top expressed markers. The cluster with the highest mCherry transcript expression was identified as the HSC cluster and utilized for further analysis.

### Split-pool barcoding scRNA-seq and data analysis

Whole kidney marrow (WKM) was isolated from each sample and dissociated into a single-cell suspension based on previously published protocol, with samples 1, 5, and 12 undergoing red blood cell lysis^116^. All other samples were interrogated with forward and side scatter (FSC/SSC) via fluorescence-activated cell sorting (FACS) to exclude erythrocytes from live WKM, following established gating strategies from previous published protocol^95^. Cells were sorted via FACS into PBS supplemented with 2% FBS. All samples were processed with a Parse WT Mini Kit according to manufacturer’s protocol for fixed cell input. Samples were processed at room temp until fixation, at which point all downstream processing occurred on ice. Libraries for split-pool barcoding scRNA-seq were prepared at the Biotechnology Center at the University of Wisconsin-Madison utilizing reagents and protocols supplied with the Parse WT Mini Kit for fixed cell input.

FASTQ files were initially processed using the “Parse Biosciences Pipeline” v.1.0.3-v.1.4.2 (Parse Biosciences) for Linux. Background was removed from unfiltered count matrices by setting a minimum transcripts per cell threshold on a per sample basis (threshold range: 332.0377 to 812.325). Further data processing was performed with R v.4.5.1, where cells with low detected features or high percent of mitochondrial reads were removed following a threshold of 2 median absolute deviations from the median, and the package “SoupX” was used to remove ambient RNA^132^. Data was normalized using size factor/log transformation and quality control was performed to remove low-quality cells and doublets. The package “batchelor” was used to reduce batch effect on reduced dimensions^133^. Clustering was performed with three levels of resolution before assigning cell types via top expressed markers. The cluster experiencing most severe batch effect after batch correction, identified as neutrophils, was removed for reclustering and further analysis. WT- or *itga4* mutant-enriched neighborhoods were defined using differential abundance analysis^134^.

### Module Score Analysis of scRNA-seq datasets

Seurat’s *AddModuleScore* function was utilized to score the transcriptomic profile of individual cells within a cluster for similarity to a curated reference gene set, termed a “module”, comparative to a stochastic set of genes selected from the aggregated expression of the cluster’s transcriptomic data as a control^135^. The number of genes to be sampled from each bin for the control module was set at 50 for each analysis. Published scRNA-seq datasets^26^ were used as curated reference gene sets for the following modules: 52-55 hpf HSPC Signature Genes (Table S4 from Xue et al.^26^), DEGs up in proliferating 55 hpf HSPCs vs non-proliferating 55 hpf HSPCs (Cluster 4 vs 0; Table S4 from Xue et al.^26^), and Cluster Identity of 55 hpf HSPCs/Progenitors (Cluster 0; Table S4 from Xue et al.^26^); and the Human Molecular Signatures Database (MSigDB) Hallmark Gene Sets for Interferon Alpha Response, Interferon Gamma Response, and Myc Targets V1^93^.

### Kidney Marrow PH3 Staining

Single cell suspension obtained from isolated kidney marrow was stained with Ghost Dye UV 450 (Cell Signaling Technology, #80862) before fixation and permeabilization. Anti-Phospho-Histone H3 (Ser 10) Alexa Fluor 647 Conjugate (Cell Signaling Technology, #3458) was used at advised concentration per cell count.

### Kidney Marrow Lineage Assessment

Isolated kidney marrow samples underwent red blood cell lysis according to established protocol^116^. Runx:mCherry+ HSPCs were interrogated with forward and side scatter (FSC/SSC) via flow cytometry to assess HSPC lineage bias, following established gating strategies from previous published protocol^95^.

### Whole-mount *in situ* hybridization (WISH)

Staining 5 dpf larvae: stage-matched larvae were fixed overnight at 4C in 4% PFA (Thermo Fisher Scientific, # J61984.AP) before proceeding with WISH based on previously published protocol^136^. Glutaraldehyde was added to a 0.2% total concentration post-fixation and larvae were bleached to remove pigmentation before permeabilizing with proteinase K. The cmyb probe was utilized as previously described to identify the presence of definitive HSPCs^137^. Imaging 5 dpf WISH larvae: Stained 5 dpf larvae were imaged with an unbiased method as previously described^138^. An 80% glycerol solution in 1x PBS buffer was utilized to mount and maintain the orientation of larvae for imaging.

### Live Imaging of Zebrafish

Larval zebrafish were mounted onto a 50 mm glass bottom dish (MatTek, #P50G-1.5-30-F) in 1% low melting point agarose (Invitrogen, #16520050) dissolved in 1x E3 (without methylene blue) plus 1x tricaine (Sigma-Aldrich, #E10521-10G). Larvae were oriented to lay laterally on the bottom of the dish, anterior on the left and posterior on the right, with tails flush against the glass. Live imaging of the tail region was performed using a Nikon AXR confocal microscope with alternating excitation of 488 nm and 568 nm laser lines. A Nikon 20x water immersion lens was used with a 2.0x zoom set within the Nikon Elements software for image capture at 40x magnification.

### Quantification of *in situ* hybridization signal

Images were analyzed unbiasedly with Fiji according to published protocol^138^. Images were inverted before converting to 8-bit. A region of interest (ROI) was manually drawn around the CHT region of the tail to measure the mean intensity value. Then, the ROI was dragged to a region in the trunk in the same fish where there was no signal from staining and the mean intensity value was recorded as a control value. The normalized mean pixel intensity value as reported is the result of the mean value from the control region subtracted from the mean value from the CHT region measured independently for each fish. Genotype was revealed after all measurements were completed.

### Surface Creation and Quantification of Volume Overlap

Images were processed with Imaris v10.2.0 (Bitplane). As an initial step, all images were cropped using the “Crop3D” function in the edit tab to confirm the z-stack was centered on Runx:mCherry+ HSPCs and to minimize autofluorescent pigment signal on the edges of the CHT. Display settings were then changed in Imaris to prevent interpolation, found in the “Display” section of the preference settings underneath the file tab. Next, the fluorescent channels were reset to baseline on the image display and “Image Proc” view was used to apply a background subtraction and gaussian filter to each channel. The filter width for background subtraction was set to 5 μm, just below the size of our cells of interest (HSPCs). The filter width for gaussian filter was 0.440 μm, the default width suggested by Imaris based on the image resolution. Once all processing steps were applied, the “Surfaces Creation Wizard” was utilized for each channel individually to render surfaces representing the fluorescent signal. The following parameters were used for the GFP channel (*zFUCCI*): “Smooth” enabled with “Surfaces Detail” set to 0.880 μm (the default suggested by Imaris based on the image resolution); the “Absolute Intensity Thresholding” method with a threshold of approximately 30 (adjusted as needed based on histogram); and, a surfaces filter for “Number of Voxels” to remove surfaces containing less than 100 voxels (equivalent to 19.36 μm^3^ based on the image resolution). The following parameters were used for the mCherry channel (*Runx:mCherry*): “Smooth” enabled with “Surfaces Detail” set to 0.880 μm (default); the “Background Subtraction (Local Contrast) Thresholding” method with “Diameter of largest sphere that fits into the object” set to 3 μm; a threshold of approximately 20 (adjusted as needed based on histogram); an “Intensity Based Split” of touching objects with the “Seed Points Diameter” set at 3 μm; a “Quality” filter for seed points according to the histogram for each replicate; and, a surfaces filter for “Number of Voxels” to remove surfaces containing less than 100 voxels (equivalent to 19.36 μm^3^ based on the image resolution). As a final curation step before identifying overlapped surface volumes, any surfaces rendered from the mCherry channel that did not match the expected morphology of Runx:mCherry+ HSPCs were manually deleted. The expected morphology was defined as round spheres of approximately 7 μm in diameter, therefore, any rendered objects displaying branching structures or long, oblique morphology were manually deleted. Upon completion of the above steps, the “Overview Statistics” were collected to record the total number of surfaces created from each channel. In the “Detailed Statistics” section of the mCherry surfaces, the “Overlapped Volume Ratio to Surfaces (Surfaces = GFP)” was selected from the “Specific Values” tab to identify each overlapping object in the image. The overlap volume ratio is reported on a scale from 0.0 to 1.0, corresponding to 0% volume overlap and 100% overlap, respectively. We defined an overlap volume ratio of 0.95-1.0 as an event indicating colocalized signal, highlighted the mCherry surfaces that met this threshold, and assigned them a unique color (white) to visually indicate colocalization. The volume ratio statistics were exported and the total number of mCherry surfaces that met the overlapped volume threshold was recorded. The percentage of mCherry surfaces with ≥95% volume overlap (out of the total number of mCherry surfaces rendered) is the final value of colocalization percentage for each replicate.

## Statistical Analysis

Prism v10.0 (GraphPad), Microsoft Excel (v2408, Microsoft Office), and the following R packages were used to perform all statistical calculations: “rstatix” (DOI:<u>10.32614/CRAN.package.rstatix</u>), “misty” (DOI:<u>10.32614/CRAN.package.misty</u>), “emmeans” (DOI:<u>10.32614/CRAN.package.emmeans</u>), and “multcomp” (DOI<u>10.32614/CRAN.package.multcomp</u>). Each analysis was chosen according to the number of conditions being compared, the number of replicates per condition, and the distribution of sample values. Statistical details for each experiment can be found in the corresponding figure legends and results section. Values of significance are summarized in this manuscript as follows: non-significant (NS) p>0.05, *p<0.05, **p<0.01, ***p<0.001, ****p<0.0001.

## Data availability

The NGS datasets generated for this study have been uploaded to the NCBI Gene Expression Omnibus (GEO) repository and can be found with the following accession numbers, as of the date of this publication:

Figure 1. Bulk RNA-seq dataset of 5 dpf Runx:mCherry+ HSPCs: GSE304735

Figure 2. scRNA-seq dataset of 5 dpf Runx:mCherry+ HSPCs: GSE304535

Figure 3. Bulk ATAC-seq dataset of 5 dpf Runx:mCherry+ HSPCs: GSE304533

Figure 5. Split-pool barcoding scRNA-seq dataset of adult WKM: GSE304975

Figure 6. Bulk ATAC-seq dataset of adult WKM: GSE329591

## Lead Contact

Requests for resources should be directed to and will be fulfilled by the lead contact, Owen J. Tamplin.

## Materials availability

Materials generated in this study are available upon request.

## Code Availability

No original code was generated for this manuscript.

## Acknowledgements

We thank the University of Wisconsin-Madison Optical Imaging Core (OIC) shared resources for their help, including use of the Nikon AXR confocal (supported by grant 1S10O34394-01) and use of Imaris (Bitplane) software. We also thank the University of Wisconsin-Madison Biotechnology Center shared resources for their help with NGS applications, including: the Gene Expression Center (GEC) for quality control and library preparation of bulk RNA-seq samples, scRNA-seq samples, and split-pool barcoding scRNA-seq samples (RRID:SCR_017757); the DNA Sequencing Core Facility for quality control of libraries prepared for bulk ATAC-seq and performing sequencing for all NGS experiments featured in this manuscript (RRID:SCR_017759); and the Bioinformatics Resource Core (BRC) for creation of a custom GRCz11 zebrafish genome that included transgenic reporters, performing alignment to the custom genome and initial quality control of sequenced reads for all NGS experiments featured in this manuscript, and Anthony Veltri for assisting in uploading all NGS datasets into GEO (RRID:SCR_017799). We thank the University of Wisconsin Carbone Cancer Center (UWCCC) shared resources for their help in usage of the Flow Cytometry Laboratory facilities and services, supported by P30 CA014520, including the following instruments: the BD FACS Aria II BSL-2 Cell Sorter, “Jill”, for many of the experiments sorting Runx:mCherry+ HSPCs (NIH Shared Instrumentation Grant #: 1S10RR025483-01, Project Title: Multi-color Benchtop Flow Cytometer); and the BD LSR Fortessa for assessing adult Runx:mCherry+ HSPC lineage bias via forward and side scatter and PH3 staining (NIH Shared Instrumentation Grant #: 1S10OD018202-01, Project Title: Special BD LSR Fortessa). We thank Dr. Michael Lasarev (University of Wisconsin-Madison Department of Biostatistics & Medical Informatics) for critical reading of the manuscript. Finally, we thank the Imaris Support team and Erich Giedzinski for guidance on analysis tools within the Imaris software. Research was supported by NIH NHLBI grants R01HL174965, R01HL142998, and R56HL142998 to OJT; NHLBI F31HL176107 to NMW; NIDDK R01DK128238 to BWB; NIDDK R01DK131162 to R.E.-P. The United States Department of Agriculture National Institute of Food and Agriculture Hatch project IOW05706 and Iowa State University start-up funds to CAC. The findings and conclusions in this publication are those of the author(s) and should not be construed to represent any official USDA or US Government determination or policy.

## Author Contributions

Conceptualization, OJT and NMW; Data curation, NMW; Formal analysis, NMW, BWB, JO, CAC; Funding acquisition, OJT and NMW; Investigation, NMW, SA, WL, BWB, JO, CAC, RC; Methodology, NMW, SA, BWB, and JO; Project administration, OJT; Resources, OJT, BWB, JO, CAC and RE-P; Software, NMW, WL, BWB, and JO; Supervision, OJT; Validation, NMW; Visualization, NMW; Writing – original draft, NMW; Writing – reviewing and editing, OJT. All authors read and approved of the final manuscript.

## References

1. Pinho S, Frenette PS. Haematopoietic stem cell activity and interactions with the niche. Nat Rev Mol Cell Biol, (2019).

2. Orkin SH, Zon LI. Hematopoiesis: an evolving paradigm for stem cell biology. Cell 132, 631–644 (2008).

3. Mikkola HKA, Orkin SH. The journey of developing hematopoietic stem cells. Development (Cambridge, England) 133, 3733–3744 (2006).

4. Hall TD, et al. Murine fetal bone marrow does not support functional hematopoietic stem and progenitor cells until birth. Nat Commun 13, 5403 (2022).

5. Ganuza M, et al. Murine foetal liver supports limited detectable expansion of life-long haematopoietic progenitors. Nat Cell Biol 24, 1475–1486 (2022).

6. Coskun S, et al. Development of the fetal bone marrow niche and regulation of HSC quiescence and homing ability by emerging osteolineage cells. Cell reports 9, 581–590 (2014).

7. Weinhaus B, et al. Differential regulation of fetal bone marrow and liver hematopoiesis by yolk-sac-derived myeloid cells. Nat Commun 16, 4427 (2025).

8. Li Y, et al. Single-Cell Analysis of Neonatal HSC Ontogeny Reveals Gradual and Uncoordinated Transcriptional Reprogramming that Begins before Birth. Cell Stem Cell 27, 732–747 e737 (2020).

9. Chen C, et al. Spatial Genome Re-organization between Fetal and Adult Hematopoietic Stem Cells. Cell Rep 29, 4200–4211 e4207 (2019).

10. Copley MR, et al. The Lin28b–let-7–Hmga2 axis determines the higher self-renewal potential of fetal haematopoietic stem cells. Nature Cell Biology 15, 916–925 (2013).

11. Jassinskaja M, et al. Comprehensive Proteomic Characterization of Ontogenic Changes in Hematopoietic Stem and Progenitor Cells. Cell Reports 21, 3285–3297 (2017).

12. McKinney-Freeman S, et al. The transcriptional landscape of hematopoietic stem cell ontogeny. Cell Stem Cell 11, 701–714 (2012).

13. Harrison DE, Zhong RK, Jordan CT, Lemischka IR, Astle CM. Relative to adult marrow, fetal liver repopulates nearly five times more effectively long-term than short-term. Exp Hematol 25, 293–297 (1997).

14. Lessard J, Faubert A, Sauvageau G. Genetic programs regulating HSC specification, maintenance and expansion. Oncogene 23, 7199–7209 (2004).

15. Manesia JK, et al. Highly proliferative primitive fetal liver hematopoietic stem cells are fueled by oxidative metabolic pathways. Stem Cell Res 15, 715–721 (2015).

16. Morrison SJ, Hemmati HD, Wandycz AM, Weissman IL. The purification and characterization of fetal liver hematopoietic stem cells. Proceedings of the National Academy of Sciences of the United States of America 92, 10302–10306 (1995).

17. Asano S, et al. MAFB regulates hematopoietic stem cell proliferation and maintenance. Sci Rep 15, 40592 (2025).

18. Kim I, Saunders TL, Morrison SJ. Sox17 dependence distinguishes the transcriptional regulation of fetal from adult hematopoietic stem cells. Cell 130, 470–483 (2007).

19. Manesia JK, et al. Distinct Molecular Signature of Murine Fetal Liver and Adult Hematopoietic Stem Cells Identify Novel Regulators of Hematopoietic Stem Cell Function. Stem Cells Dev 26, 573–584 (2017).

20. Popescu DM, et al. Decoding human fetal liver haematopoiesis. Nature 574, 365–371 (2019).

21. Gao S, Liu F. Fetal liver: an ideal niche for hematopoietic stem cell expansion. Sci China Life Sci 61, 885–892 (2018).

22. Mahony CB, Bertrand JY. How HSCs Colonize and Expand in the Fetal Niche of the Vertebrate Embryo: An Evolutionary Perspective. Front Cell Dev Biol 7, 34 (2019).

23. Agrawal H, Mehatre SH, Khurana S. The hematopoietic stem cell expansion niche in fetal liver: Current state of the art and the way forward. Exp Hematol 136, 104585 (2024).

24. Abrams EW, Mullins MC. Early zebrafish development: it’s in the maternal genes. Current Opinion in Genetics &amp; Development 19, 396–403 (2009).

25. Aharon D, Marlow FL. Sexual determination in zebrafish. Cell Mol Life Sci 79, 8 (2021).

26. Xue Y, et al. A 3D Atlas of Hematopoietic Stem and Progenitor Cell Expansion by Multi-dimensional RNA-Seq Analysis. Cell Rep 27, 1567–1578 e1565 (2019).

27. Calvanese V, et al. Mapping human haematopoietic stem cells from haemogenic endothelium to birth. Nature 604, 534–540 (2022).

28. Guo XL, et al. Fetal hepatocytes protect the HSPC genome via fetuin-A. Nature 637, 402–411 (2025).

29. Agarwala S, et al. Defining the ultrastructure of the hematopoietic stem cell niche by correlative light and electron microscopy. Elife 11, (2022).

30. Tamplin OJ, et al. Hematopoietic Stem Cell Arrival Triggers Dynamic Remodeling of the Perivascular Niche. Cell 160, 241–252 (2015).

31. Kissa K, Herbomel P. Blood stem cells emerge from aortic endothelium by a novel type of cell transition. Nature 464, 112–115 (2010).

32. Bertrand JY, Chi NC, Santoso B, Teng S, Stainier DYR, Traver D. Haematopoietic stem cells derive directly from aortic endothelium during development. Nature 464, 108–111 (2010).

33. He Q, et al. Inflammatory signaling regulates hematopoietic stem and progenitor cell emergence in vertebrates. Blood 125, 1098–1106 (2015).

34. Espín-Palazón R, et al. Proinflammatory signaling regulates hematopoietic stem cell emergence. Cell 159, 1070–1085 (2014).

35. Campbell CA, et al. p65 signaling dynamics drive the developmental progression of hematopoietic stem and progenitor cells through cell cycle regulation. Nat Commun 15, 7787 (2024).

36. Li Y, et al. Inflammatory signaling regulates embryonic hematopoietic stem and progenitor cell production. Genes &amp; Development 28, 2597–2612 (2014).

37. Sawamiphak S, Kontarakis Z, Stainier Didier YR. Interferon Gamma Signaling Positively Regulates Hematopoietic Stem Cell Emergence. Developmental Cell 31, 640–653 (2014).

38. Cheng X, et al. Nod1-dependent NF-kB activation initiates hematopoietic stem cell specification in response to small Rho GTPases. Nat Commun 14, 7668 (2023).

39. Frame JM, et al. Metabolic Regulation of Inflammasome Activity Controls Embryonic Hematopoietic Stem and Progenitor Cell Production. Dev Cell 55, 133–149.e136 (2020).

40. Lefkopoulos S, et al. Repetitive Elements Trigger RIG-I-like Receptor Signaling that Regulates the Emergence of Hematopoietic Stem and Progenitor Cells. Immunity 53, 934–951 e939 (2020).

41. Thambyrajah R, et al. IκBα controls dormancy in hematopoietic stem cells via retinoic acid during embryonic development. Nat Commun 15, 4673 (2024).

42. Mariani SA, et al. Pro-inflammatory aorta-associated macrophages are involved in embryonic development of hematopoietic stem cells. Immunity 50, 1439–1452.E1435 (2019).

43. Tie R, et al. Interleukin-6 signaling regulates hematopoietic stem cell emergence. Experimental & Molecular Medicine 51, 1–12 (2019).

44. Murayama E, et al. Tracing Hematopoietic Precursor Migration to Successive Hematopoietic Organs during Zebrafish Development. Immunity 25, 963–975 (2006).

45. Ganuza M, et al. Murine hemogenic endothelial precursors display heterogeneous hematopoietic potential ex vivo. Exp Hematol 51, 25–35 e26 (2017).

46. Chen MJ, Yokomizo T, Zeigler BM, Dzierzak E, Speck NA. Runx1 is required for the endothelial to haematopoietic cell transition but not thereafter. Nature 457, 887–891 (2009).

47. Chen MJ, et al. Erythroid/myeloid progenitors and hematopoietic stem cells originate from distinct populations of endothelial cells. Cell stem cell 9, 541–552 (2011).

48. Ulloa BA, et al. Definitive hematopoietic stem cells minimally contribute to embryonic hematopoiesis. Cell Rep 36, 109703 (2021).

49. Henninger J, et al. Clonal fate mapping quantifies the number of haematopoietic stem cells that arise during development. Nat Cell Biol 19, 17–27 (2017).

50. Wattrus SJ, et al. Quality assurance of hematopoietic stem cells by macrophages determines stem cell clonality. Science 377, 1413–1419 (2022).

51. Krenn PW, Montanez E, Costell M, Fassler R. Integrins, anchors and signal transducers of hematopoietic stem cells during development and in adulthood. Curr Top Dev Biol 149, 203–261 (2022).

52. Li D, et al. VCAM-1+ macrophages guide the homing of HSPCs to a vascular niche. Nature 564, 119–124 (2018).

53. Priestley GV, Scott LM, Ulyanova T, Papayannopoulou T. Lack of alpha4 integrin expression in stem cells restricts competitive function and self-renewal activity. Blood 107, 2959–2967 (2006).

54. Qian H, et al. Distinct roles of integrins alpha6 and alpha4 in homing of fetal liver hematopoietic stem and progenitor cells. Blood 110, 2399–2407 (2007).

55. Gribi R, Hook L, Ure J, Medvinsky A. The differentiation program of embryonic definitive hematopoietic stem cells is largely alpha4 integrin independent. Blood 108, 501–509 (2006).

56. Williams DA, Rios M, Stephens C, Patel VP. Fibronectin and VLA-4 in haematopoietic stem cell–microenvironment interactions. Nature 352, 438–441 (1991).

57. Potocnik AJ, Brakebusch C, Fässler R. Fetal and adult hematopoietic stem cells require beta1 integrin function for colonizing fetal liver, spleen, and bone marrow. Immunity 12, 653–663 (2000).

58. Hirsch E, Iglesias A, Potocnik AJ, Hartmann U, Fässler R. Impaired migration but not differentiation of haematopoietic stem cells in the absence of β1 integrins. Nature 380, 171–175 (1996).

59. Arroyo AG, Yang JT, Rayburn H, Hynes RO. Differential requirements for alpha4 integrins during fetal and adult hematopoiesis. Cell 85, 997–1008 (1996).

60. Arroyo AG, Yang JT, Rayburn H, Hynes RO. Alpha4 integrins regulate the proliferation/differentiation balance of multilineage hematopoietic progenitors in vivo. Immunity 11, 555–566 (1999).

61. Elices MJ, et al. VCAM-1 on activated endothelium interacts with the leukocyte integrin VLA-4 at a site distinct from the VLA-4/fibronectin binding site. Cell 60, 577–584 (1990).

62. Bornhorst D, et al. The heart is a resident tissue for hematopoietic stem and progenitor cells in zebrafish. Nat Commun 15, 7589 (2024).

63. Santis Larrain O, et al. Vcam1 in endothelial and stromal cells regulates hematopoietic stem cell contact with the niche. Development, (2026).

64. Dann E, Henderson NC, Teichmann SA, Morgan MD, Marioni JC. Differential abundance testing on single-cell data using k-nearest neighbor graphs. Nat Biotechnol 40, 245–253 (2022).

65. Essers MA, et al. IFNalpha activates dormant haematopoietic stem cells in vivo. Nature 458, 904–908 (2009).

66. Ueda Y, Cain DW, Kuraoka M, Kondo M, Kelsoe G. IL-1R type I-dependent hemopoietic stem cell proliferation is necessary for inflammatory granulopoiesis and reactive neutrophilia. J Immunol 182, 6477–6484 (2009).

67. Baldridge MT, King KY, Boles NC, Weksberg DC, Goodell MA. Quiescent haematopoietic stem cells are activated by IFN-gamma in response to chronic infection. Nature 465, 793–797 (2010).

68. Pietras EM, et al. Re-entry into quiescence protects hematopoietic stem cells from the killing effect of chronic exposure to type I interferons. The Journal of experimental medicine 211, 245–262 (2014).

69. Saka S, et al. Chromatin factor YY1 controls fetal hematopoietic stem cell migration and engraftment in mice. J Clin Invest 135, (2025).

70. Lu Z, et al. Polycomb Group Protein YY1 Is an Essential Regulator of Hematopoietic Stem Cell Quiescence. Cell Rep 22, 1545–1559 (2018).

71. Ito K, et al. A PML-PPAR-delta pathway for fatty acid oxidation regulates hematopoietic stem cell maintenance. Nat Med 18, 1350–1358 (2012).

72. Xue Y, Gao S, Liu F. Genome-wide analysis of the zebrafish Klf family identifies two genes important for erythroid maturation. Dev Biol 403, 115–127 (2015).

73. Xue Y, Lv J, Zhang C, Wang L, Ma D, Liu F. The Vascular Niche Regulates Hematopoietic Stem and Progenitor Cell Lodgment and Expansion via klf6a-ccl25b. Developmental Cell, (2017).

74. Matsumoto N, et al. Developmental regulation of yolk sac hematopoiesis by Kruppel-like factor 6. Blood 107, 1357–1365 (2006).

75. Collins A, et al. Maternal inflammation regulates fetal emergency myelopoiesis. Cell 187, 1402–1421.e1421 (2024).

76. Chia JJ, et al. Distinct roles for NF-kappaB in hematopoietic stem cells and the bone marrow milieu in promoting hematopoietic aging. Cell Rep 44, 116193 (2025).

77. Morino-Koga S, Yokomizo T. Deciphering hematopoietic stem cell development: key signaling pathways and mechanisms. Frontiers in Cell and Developmental Biology Volume 12 - 2024, (2024).

78. Antonchuk J, Sauvageau G, Humphries RK. HOXB4 overexpression mediates very rapid stem cell regeneration and competitive hematopoietic repopulation. Exp Hematol 29, 1125–1134 (2001).

79. Heinz S, et al. Simple combinations of lineage-determining transcription factors prime cis-regulatory elements required for macrophage and B cell identities. Mol Cell 38, 576–589 (2010).

80. Cabezas-Wallscheid N, et al. Vitamin A-Retinoic Acid Signaling Regulates Hematopoietic Stem Cell Dormancy. Cell, (2017).

81. Rubin JD, et al. Transcription factor enrichment analysis (TFEA) quantifies the activity of multiple transcription factors from a single experiment. Commun Biol 4, 661 (2021).

82. Bentsen M, et al. ATAC-seq footprinting unravels kinetics of transcription factor binding during zygotic genome activation. Nature Communications 11, 4267 (2020).

83. Edginton-White B, et al. A genome-wide relay of signalling-responsive enhancers drives hematopoietic specification. Nat Commun 14, 267 (2023).

84. Obier N, et al. Cooperative binding of AP-1 and TEAD4 modulates the balance between vascular smooth muscle and hemogenic cell fate. Development 143, 4324–4340 (2016).

85. Sugiyama M, et al. Illuminating cell-cycle progression in the developing zebrafish embryo. Proc Natl Acad Sci U S A 106, 20812–20817 (2009).

86. Lin H-F, et al. Analysis of thrombocyte development in CD41-GFP transgenic zebrafish. Blood 106, 3803–3810 (2005).

87. Hyduk SJ, Oh J, Xiao H, Chen M, Cybulsky MI. Paxillin selectively associates with constitutive and chemoattractant-induced high-affinity alpha4beta1 integrins: implications for integrin signaling. Blood 104, 2818–2824 (2004).

88. Kanda E, Jin ZH, Mizuchi D, Arai A, Miura O. Activation of Rac and tyrosine phosphorylation of cytokine receptors induced by cross-linking of integrin alpha4beta1 and cell adhesion in hematopoietic cells. Biochem Biophys Res Commun 301, 934–940 (2003).

89. Gao X, Zhang J, Tamplin OJ. The aging hematopoietic stem cell niche: a mini review. Front Hematol 4, (2025).

90. Tang Q, et al. Dissecting hematopoietic and renal cell heterogeneity in adult zebrafish at single-cell resolution using RNA sequencing. The Journal of experimental medicine 214, 2875–2887 (2017).

91. Han X, Pang K, Liu X, Zhou J, Zhu J, Yuan H. Novel roles for CREG1 in hematopoiesis revealed by single-cell RNA sequencing. Cell Biosci 15, 74 (2025).

92. Geiger H, Denkinger M, Schirmbeck R. Hematopoietic stem cell aging. Curr Opin Immunol 29, 86–92 (2014).

93. Liberzon A, Birger C, Thorvaldsdottir H, Ghandi M, Mesirov JP, Tamayo P. The Molecular Signatures Database (MSigDB) hallmark gene set collection. Cell Syst 1, 417–425 (2015).

94. Tanaka-Yano M, et al. Dynamic activity of Erg promotes maturation of the hematopoietic system. Blood Adv 10, 759–773 (2026).

95. Traver D, Paw BH, Poss KD, Penberthy WT, Lin S, Zon LI. Transplantation and in vivo imaging of multilineage engraftment in zebrafish bloodless mutants. Nature Immunology 4, 1238–1246 (2003).

96. Müller AM, Medvinsky A, Strouboulis J, Grosveld F, Dzierzak E. Development of hematopoietic stem cell activity in the mouse embryo. Immunity 1, 291–301 (1994).

97. Bogeska R, et al. Inflammatory exposure drives long-lived impairment of hematopoietic stem cell self-renewal activity and accelerated aging. Cell Stem Cell 29, 1273–1284.e1278 (2022).

98. Rybtsov S, Ivanovs A, Zhao S, Medvinsky A. Concealed expansion of immature precursors underpins acute burst of adult HSC activity in foetal liver. Development (Cambridge, England) 143, 1284–1289 (2016).

99. Caiado F, Kovtonyuk LV, Gonullu NG, Fullin J, Boettcher S, Manz MG. Aging drives Tet2+/- clonal hematopoiesis via IL-1 signaling. Blood 141, 886–903 (2023).

100. Su TY, et al. Aging is associated with functional and molecular changes in distinct hematopoietic stem cell subsets. Nature Communications 15, (2024).

101. Colom Diaz PA, Mistry JJ, Trowbridge JJ. Hematopoietic stem cell aging and leukemia transformation. Blood 142, 533–542 (2023).

102. Chambers SM, Shaw CA, Gatza C, Fisk CJ, Donehower LA, Goodell MA. Aging hematopoietic stem cells decline in function and exhibit epigenetic dysregulation. PLoS Biol 5, e201 (2007).

103. Xia J, et al. Activation of lineage competence in hemogenic endothelium precedes the formation of hematopoietic stem cell heterogeneity. Cell Res 33, 448–463 (2023).

104. Azzoni E, et al. The onset of circulation triggers a metabolic switch required for endothelial to hematopoietic transition. Cell Rep 37, 110103 (2021).

105. Pv A, Mehatre SH, Verfaillie CM, Alam MT, Khurana S. Glycolytic state of aortic endothelium favors hematopoietic transition during the emergence of definitive hematopoiesis. Sci Adv 10, eadh8478 (2024).

106. Gerri C, Marass M, Rossi A, Stainier DYR. Hif-1α and Hif-2α regulate hemogenic endothelium and hematopoietic stem cell formation in zebrafish. Blood 131, 963–973 (2018).

107. Nakamura-Ishizu A, Ito K, Suda T. Hematopoietic Stem Cell Metabolism during Development and Aging. Dev Cell 54, 239–255 (2020).

108. Ansó E, et al. The mitochondrial respiratory chain is essential for haematopoietic stem cell function. Nat Cell Biol 19, 614–625 (2017).

109. Scott LM, Priestley GV, Papayannopoulou T. Deletion of alpha4 integrins from adult hematopoietic cells reveals roles in homeostasis, regeneration, and homing. Molecular and Cellular Biology 23, 9349–9360 (2003).

110. Ramirez P, et al. BIO5192, a small molecule inhibitor of VLA-4, mobilizes hematopoietic stem and progenitor cells. Blood 114, 1340–1343 (2009).

111. Khurana S, et al. Outside-in integrin signalling regulates haematopoietic stem cell function via Periostin-Itgav axis. Nat Commun 7, 13500 (2016).

112. Percie du Sert N, et al. The ARRIVE guidelines 2.0: Updated guidelines for reporting animal research. PLoS Biol 18, e3000410 (2020).

113. Westerfield M. The Zebrafish Book. A Guide for the Laboratory Use of Zebrafish (Danio rerio), 4th edn. University of Oregon Press (2000).

114. King AC, Gut M, Zenker AK. Shedding new light on early sex determination in zebrafish. Arch Toxicol 94, 4143–4158 (2020).

115. Bresciani E, Broadbridge E, Liu PP. An efficient dissociation protocol for generation of single cell suspension from zebrafish embryos and larvae. MethodsX 5, 1287–1290 (2018).

116. Fraint E, Lv P, Liu F, Bowman TV, Tamplin OJ. Hematopoietic Stem and Progenitor Cell Identification and Transplantation in Zebrafish. Methods Mol Biol 2567, 233–249 (2023).

117. Jiang H, Lei R, Ding SW, Zhu S. Skewer: a fast and accurate adapter trimmer for next-generation sequencing paired-end reads. BMC Bioinformatics 15, 182 (2014).

118. Dobin A, et al. STAR: ultrafast universal RNA-seq aligner. Bioinformatics 29, 15–21 (2013).

119. Li B, Dewey CN. RSEM: accurate transcript quantification from RNA-Seq data with or without a reference genome. BMC Bioinformatics 12, 323 (2011).

120. Robinson MD, Oshlack A. A scaling normalization method for differential expression analysis of RNA-seq data. Genome Biol 11, R25 (2010).

121. Reiner A, Yekutieli D, Benjamini Y. Identifying differentially expressed genes using false discovery rate controlling procedures. Bioinformatics 19, 368–375 (2003).

122. Aken BL, et al. The Ensembl gene annotation system. Database (Oxford) 2016, (2016).

123. Subramanian A, et al. Gene set enrichment analysis: a knowledge-based approach for interpreting genome-wide expression profiles. Proc Natl Acad Sci U S A 102, 15545–15550 (2005).

124. Buenrostro JD, Wu B, Chang HY, Greenleaf WJ. ATAC-seq: A Method for Assaying Chromatin Accessibility Genome-Wide. Curr Protoc Mol Biol 109, 21.29.21-21.29.29 (2015).

125. Buenrostro JD, et al. Single-cell chromatin accessibility reveals principles of regulatory variation. Nature 523, 486–490 (2015).

126. Ross-Innes CS, et al. Differential oestrogen receptor binding is associated with clinical outcome in breast cancer. Nature 481, 389–393 (2012).

127. Zhu LJ, et al. ChIPpeakAnno: a Bioconductor package to annotate ChIP-seq and ChIP-chip data. BMC Bioinformatics 11, 237 (2010).

128. Ritchie ME, et al. limma powers differential expression analyses for RNA-sequencing and microarray studies. Nucleic Acids Res 43, e47 (2015).

129. Bentsen M, et al. ATAC-seq footprinting unravels kinetics of transcription factor binding during zygotic genome activation. Nat Commun 11, 4267 (2020).

130. Quinlan AR, Hall IM. BEDTools: a flexible suite of utilities for comparing genomic features. Bioinformatics 26, 841–842 (2010).

131. Ovek Baydar D, et al. JASPAR 2026: expansion of transcription factor binding profiles and integration of deep learning models. Nucleic Acids Res 54, D184–d193 (2026).

132. Young MD, Behjati S. SoupX removes ambient RNA contamination from droplet-based single-cell RNA sequencing data. Gigascience 9, (2020).

133. Haghverdi L, Lun ATL, Morgan MD, Marioni JC. Batch effects in single-cell RNA-sequencing data are corrected by matching mutual nearest neighbors. Nat Biotechnol 36, 421–427 (2018).

134. Zhao J, et al. Detection of differentially abundant cell subpopulations in scRNA-seq data. Proc Natl Acad Sci U S A 118, (2021).

135. Tirosh I, et al. Dissecting the multicellular ecosystem of metastatic melanoma by single-cell RNA-seq. Science 352, 189–196 (2016).

136. Thisse C, Thisse B. High-resolution in situ hybridization to whole-mount zebrafish embryos. Nature Protocols 3, 59–69 (2008).

137. Liao EC, Paw BH, Oates AC, Pratt SJ, Postlethwait JH, Zon LI. SCL/Tal-1 transcription factor acts downstream of cloche to specify hematopoietic and vascular progenitors in zebrafish. Genes &amp; Development 12, 621–626 (1998).

138. Dobrzycki T, Krecsmarik M, Monteiro R. Genotyping and Quantification of In Situ Hybridization Staining in Zebrafish. Journal of visualized experiments : JoVE, (2020).

139. Hothorn T, Bretz F, Westfall P. Simultaneous inference in general parametric models. Biom J 50, 346–363 (2008).

