## Supplementary Figures and Methods for "Niche contact during development programs hematopoietic stem cells"

**Description of Additional Supplementary Files**

**Supplementary Methods**

**Supplementary Figures 1-9**

**Supplementary Tables 1-5**

**Table S1.** DGE, GO, and GSEA results from bulk RNA-seq of 5 dpf sorted Runx:mCherry+ HSPCs (*itga4*$^{UWT013}$ vs WT). Related to Main Figure 1.

**Table S2.** DGE and GSEA results from scRNA-seq of 5 dpf sorted Runx:mCherry+ HSPCs (*itga4*$^{UWT013}$ vs WT). Related to Main Figure 2 and Supplemental Figure S2.

**Table S3.** Diffbind, HOMER motif analysis, TFEA, TOBIAS footprinting, and GO analysis results from bulk ATAC-seq of 5 dpf sorted Runx:mCherry+ HSPCs. Related to Main Figure 3 and Supplemental Figure S3.

**Table S4.** Pairwise DGE analyses from split-pool barcoding scRNA-seq of 6 mpf *itga4*$^{UWT013}$, 6 mpf *vcam1b*$^{UWT007}$, 6 mpf WT, and 24 mpf WT WKM (red blood cells excluded). Related to Main Figure 5.

**Table S5.** Diffbind, HOMER motif analysis, and TOBIAS footprinting analysis results from bulk ATAC-seq of sorted Runx:mCherry+ HSPCs from 6 mpf *itga4*$^{UWT013}$, 6 mpf *vcam1b*$^{UWT007}$, 6 mpf WT, and 24 mpf WT WKM. Related to Main Figure 6.

### Supplementary Methods

#### Sequencing and Analysis

##### Bulk RNA-seq and data analysis

Runx:mCherry+ HSPCs were sorted via FACS directly into Trizol LS (Thermo Fisher Scientific, #10296028) and stored at -80 °C until all replicates were collected. Once all replicates were collected, samples were brought to room temperature and the protocol for total RNA isolation was completed based on Invitrogen protocol Cat. No. 10296-010. Isolated RNA was sent to the Biotechnology Center at the University of Wisconsin-Madison for quality control analysis via TapeStation, library preparation, and sequencing. RNA libraries for RNA-seq were prepared using Takara SMART-Seq v4 Ultra Low Input RNA/NexteraXT (polyA enrichment) following manufacturer's protocols.

Reads underwent adapter trimming with “skewer” v.0.2.2b (PMID:24925680), then were mapped to a custom GRCz11 reference genome with “STAR” v.2.5.3a (PMID: 23104886). The GRCz11 reference genome was customized to include inserted transgenic reporter constructs (e.g. *Runx:mCherry*). Mapped reads were quantified with “RSEM” v.1.2.31 (PMID:21816040) and processed via “edgeR” v.3.34.0 (PMID:19910308) to independently filter samples with the *filterByExpr* function to remove genes with less than 10 read counts, normalize each sample by the method of trimmed mean of M-values (PMID:20196867), and perform analysis of differentially expressed genes between samples while controlling the false discovery rate (PMID:12584122). Finally, transcripts were annotated with the “Ensembl” annotation system (PMID:27337980) and DEGs underwent downstream analysis with GSEA (PMID:16199517).

**Bulk ATAC-seq and data analysis**

Runx:mCherry+ HSPCs were sorted via FACS directly into PBS supplemented with 2% FBS for downstream processing. Libraries for ATAC-seq were prepared based off previously published protocol for ATAC-seq (PMID:25559105). Primer sequences used to generate the libraries were referenced from previously published protocol (PMID:26083756), with additional size selection of fragments and library cleanup using AmPure XP Beads (Beckman Coulter, #A63880). The number of PCR cycles for library amplification was optimized for organism and cell type, which deviated from the referenced published protocols. Unpooled libraries were sent to the Biotechnology Center at the University of Wisconsin-Madison for quality control analysis via TapeStation and sequencing.

Sequences were analyzed by ENCODE ATAC-seq pipeline (v2.2.3) with customized GRCz11 genome with additional DsRed2, ZsGreen, mCherry, and EGFP gene sequences. The conservative set selected by IDR (Irreproducible Discovery Rate) were used for Downstream analysis. DiffBind v3.16.0 (DOI:10.18129) was used to call differential accessible sites (DAS). ATAC-Seq peaks were annotated nearest gene symbols by ChIPpeakAnno v3.40.0 (DOI:10.1186). The motif enrichment analysis was performed by TFEA v1.1.2 (DOI:10.1038). The motifs enrichment was also performed for the DAS filtered with FoldChange>1.1 and P.value<0.05 by Homer v5.1 (PMID:20513432). The promoter and gene body usage analysis were performed by LIMMA v3.62.2 (DOI:10.1093) voom for the reads counts in each region. The promoters were defined as upstream 1K to downstream 500bp from the TSS. The gene body was defined as gene ranges without the promoter.

Transcription factor (TF) footprinting analysis was performed using TOBIAS v0.17.3 (PMID:32848148). To generate a common reference set of accessible chromatin regions for footprinting analysis, the IDR optimal peak sets from the wild-type and mutant samples were combined, sorted, and merged using bedtools v2.31.1 (PMID:20110278) merge to create a consensus peak set. Peak annotations and genomic regions of interest were generated using the TOBIAS region preparation utilities with the parameters --feature CDS, --feature_anchor start, and --distance 20000 10000, defining regions spanning 20 kb upstream and 10 kb downstream of coding sequence start sites. TF binding motifs were obtained from the JASPAR2026 CORE (PMID:41325984) non-redundant position frequency matrix collection. TOBIAS ATACorrect was used to correct Tn5 transposase sequence bias for each sample, followed by FootprintScores to calculate genome-wide footprint scores. Differential TF binding between wild-type and mutant samples was assessed using BINDetect, which integrates corrected footprint scores with motif occurrences to identify TFs with differential predicted binding occupancy between the two conditions.

**scRNA-seq and data analysis**

Runx:mCherry+ HSPCs were sorted via FACS directly into 0.2mL PCR strip tubes with PBS supplemented with 2% FBS for a “sort and load” protocol directly into the library preparation stage. Samples were then immediately loaded onto a 10X Genomics 3' v3.1 chip for processing according to manufacturer's instructions. Libraries for scRNA-seq were prepared at the Biotechnology Center at the University of Wisconsin-Madison utilizing reagents and protocols supplied with the 10X Genomics 3' v3.1 kit under standard conditions.

The “cell ranger” v.9.0.0 pipeline (10X Genomics) was utilized for initial processing and alignment to the custom GRCz11 reference genome. Further data processing was completed via R v.4.5.1, where cells with low detected features or high percent of mitochondrial reads were removed by a threshold of 2 median absolute deviations from the median and the package “SoupX” was used to remove ambient RNA (PMID:33367645). Data was normalized using size factor/log transformation and quality control was performed to remove low quality cells and doublets. The package “batchelor” was used to reduce batch effect on reduced dimensions (PMID:29608177). Clustering was performed with three levels of resolution before assigning cell types via top expressed markers. The cluster with the highest mCherry transcript expression was identified as the HSC cluster and utilized for further analysis. WT- or *itga4* mutant-enriched neighborhoods were defined using differential abundance analysis (DOI:10.1101/711929).

**Module Score Analysis of scRNA-seq datasets**

Seurat's *AddModuleScore* function was utilized to score the transcriptomic profile of individual cells within a cluster for similarity to a curated reference gene set, termed a "module", comparative to a stochastic set of genes selected from the aggregated expression of the cluster's transcriptomic data as a control (PMID:27124452). The number of genes to be sampled from each bin for the control module was set at 50 for each analysis. Published scRNA-seq datasets were used as curated reference gene sets for the following modules: 52-55 hpf HSPC Signature Genes, DEGs up in proliferating 55 hpf HSPCs vs non-proliferating 55 hpf HSPCs, and Cluster Identity of 55 hpf HSPCs/Progenitors (PMID:31042481); and the Human Molecular Signatures Database (MSigDB) Hallmark Gene Sets for Interferon Alpha Response, Interferon Gamma Response, and Myc Targets V1 (PMID:26771021).

**Supplementary Figures**

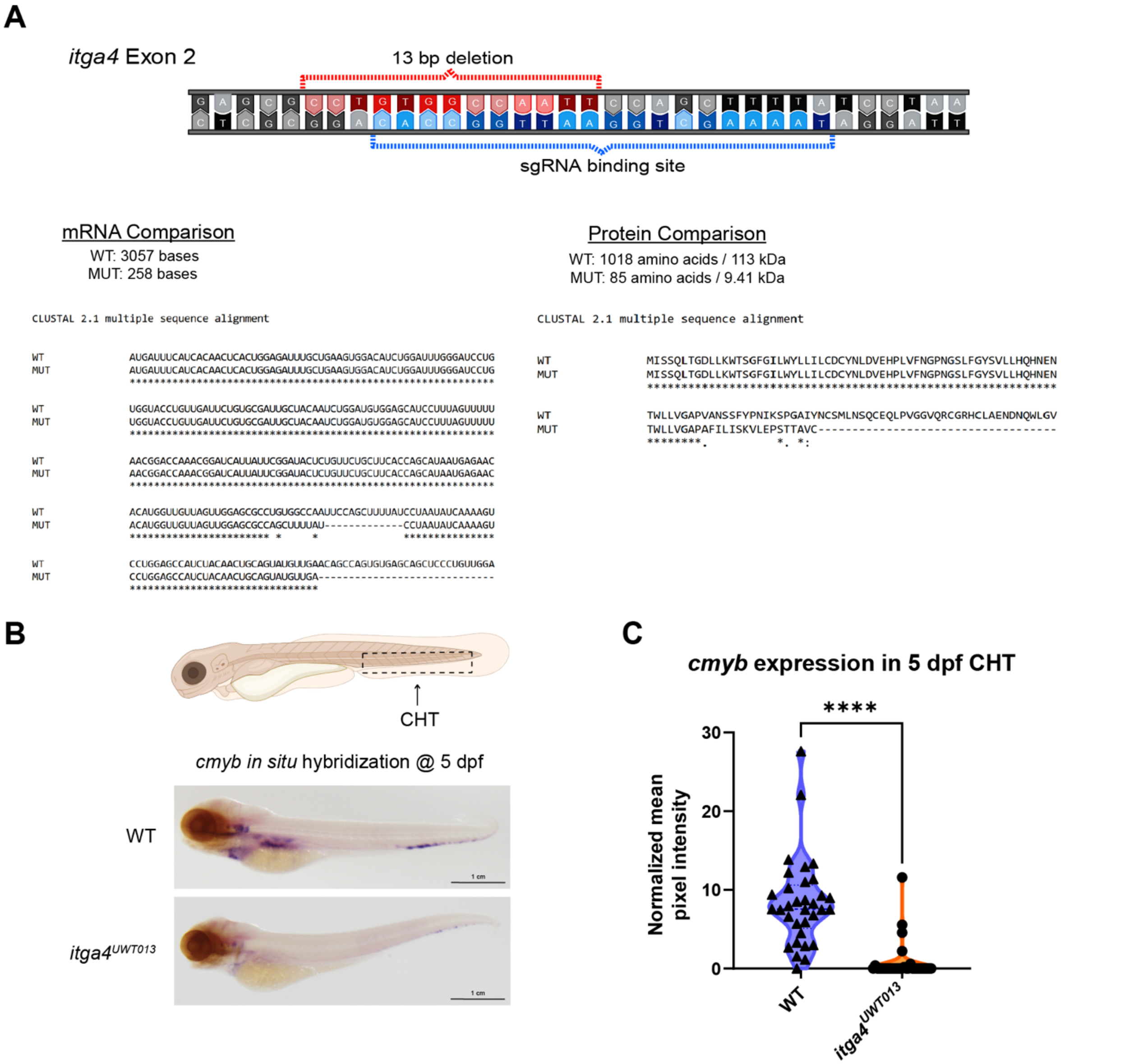


**Supplementary Figure 1. Creation and Characterization of *itga4*$^{UWT013}$ zebrafish line. Related to Main Figure 1.**

**A)** Diagram at the top, center, represents the binding site recognized by the sgRNA that was used to create the *itga4*$^{UWT013}$ line, along with the resulting 13 base pair (bp) deletion resulting from successful CRISPR-mediated mutation (created with Biorender). This occurs in exon 2 of the *itga4* gene. A comparison between WT and homozygous mutant (MUT) mRNA, bottom left, indicates significant truncation of the resulting transcribed product (3057 base length in WT vs 258 base length in MUT). Sequence alignment with CLUSTAL 2.1 demonstrates remaining conservation in the mRNA products (* = conserved identical sequence, - = gap, no symbol = non-conservative mutation). A comparison between WT and MUT protein, bottom right, similarly shows significant truncation of the translated MUT protein product, as expected (1018 amino acids & 113 kDa predicted in WT vs 85 amino acids & 9.41 kDa predicted in MUT). Sequence alignment with CLUSTAL 2.1 demonstrates remaining conservation in the protein products (* = conserved identical sequence, : = conservative mutation, . = semi-conservative mutation, - = gap, no symbol = non-conservative mutation). **B)** Representative *in situ* hybridization images probing for *cmyb* at 5 dpf in each condition. Dark purple stains indicate positive expression. Representative figure of larval zebrafish at the top indicates the region of interest, the CHT, marked with an arrow and a box with dashed lines. **C)** Quantification of positive staining for *cmyb* via *in situ* hybridization. Values were normalized to a chosen control region within the trunk of each zebrafish that was negative for *cmyb* expression. Each symbol in a violin plot represents one larval fish (n=33 for WT and n=33 for $itga4^{UWT013}$, replicates were processed in one experiment and imaged on a separate experimental day). Statistical analysis via unpaired Mann-Whitney test (****p<0.0001); WT 5 dpf median= 7.56 and IQR= 5.50; *itga4* mutant 5 dpf median= 0.00 and IQR= 0.00.

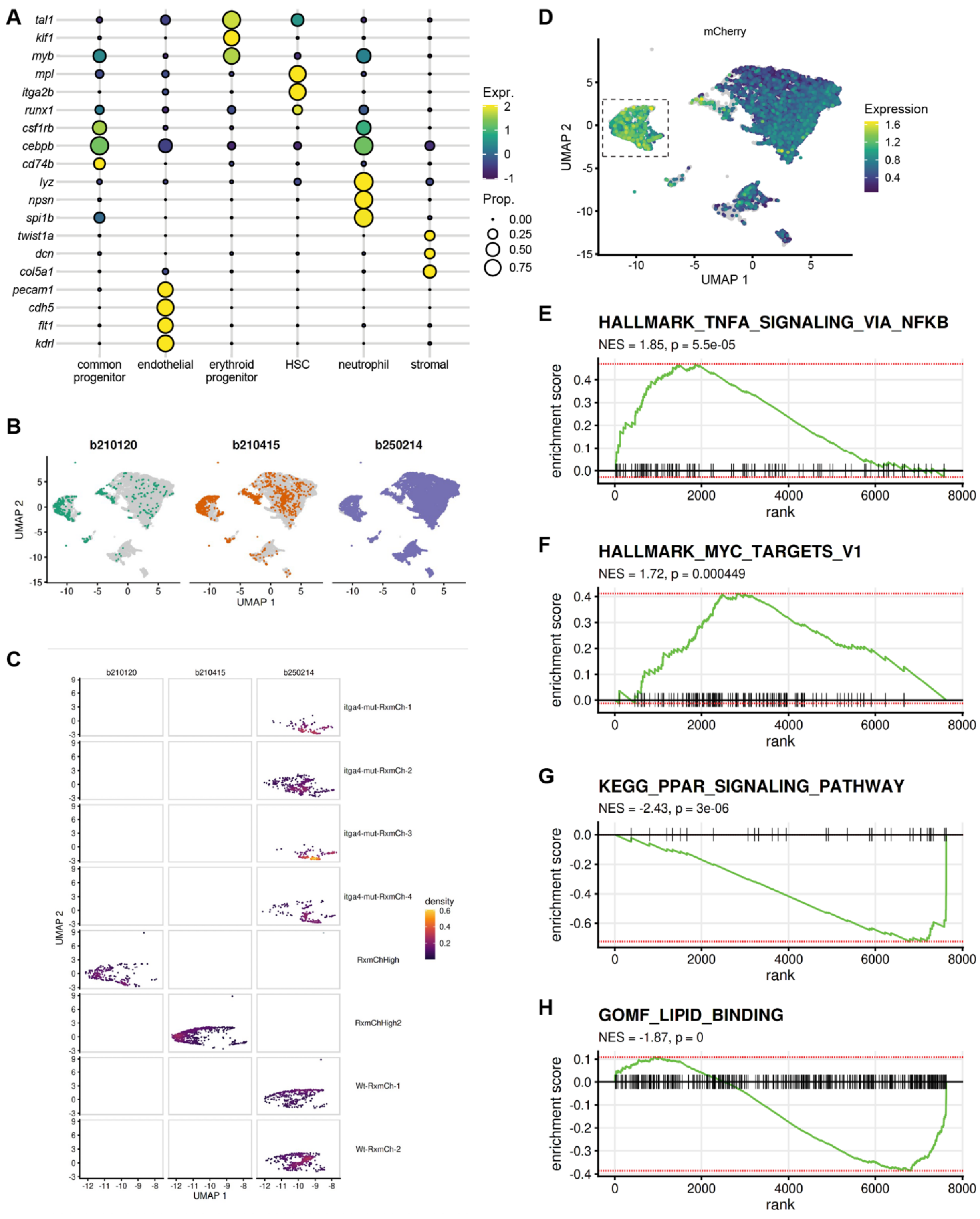


**Supplementary Figure 2. scRNA-seq of 5 dpf Runx:mCherry+ HSPCs sorted from pooled *itga4*$^{UWT013}$ and WT larvae. Related to Main Figure 2.**

**A)** Bubble plot showing differential expression of individual genes used to determine cluster identities (see **Fig. 2A**). **B)** A comparison of UMAP projections of all sequenced cells after batch correction, separated into individual projections based on sequencing date (labeled at top and color coded: b210120 = green, left; b210415 = orange, center; b250214 = purple, right. **C)** A comparison of UMAP projections of the HSC cluster after batch correction, separated by sample condition (labeled along y-coordinate, right side) and date of sequencing (labeled along x-coordinate, top, with “b” standing for “batch” and date written as year-month-day). Note that “RxmChHigh” and “RxmChHigh2” are WT conditions. The sample “itga4-mut-RxmCh-2” was split into two samples before library prep, making the sample “itga4-mut-RxmCh-3” a technical replicate. **D)** Expression of *mCherry* transcript is high in HSC cluster, consistent with these being *Runx:mCherry*$^{High}$ cells. The HSC cluster (dashed box) was used for further analysis (see **Fig. 2C-G**). **E-H)** Top GSEA pathways represented by DEGs between *itga4*$^{UWT013}$ and WT HSCs (panel **C and Table S2**); **E)** TNFA_SIGNALING_VIA_NFκB; **F)** MYC_TARGETS_V1; **G)** PPAR_SIGNALING_PATHWAY; **H)** LIPID_BINDING; positive NES is upregulated in *itga4*$^{UWT013}$ HSCs compared to WT and negative NES is upregulated in WT HSCs compared to *itga4*$^{UWT013}$.

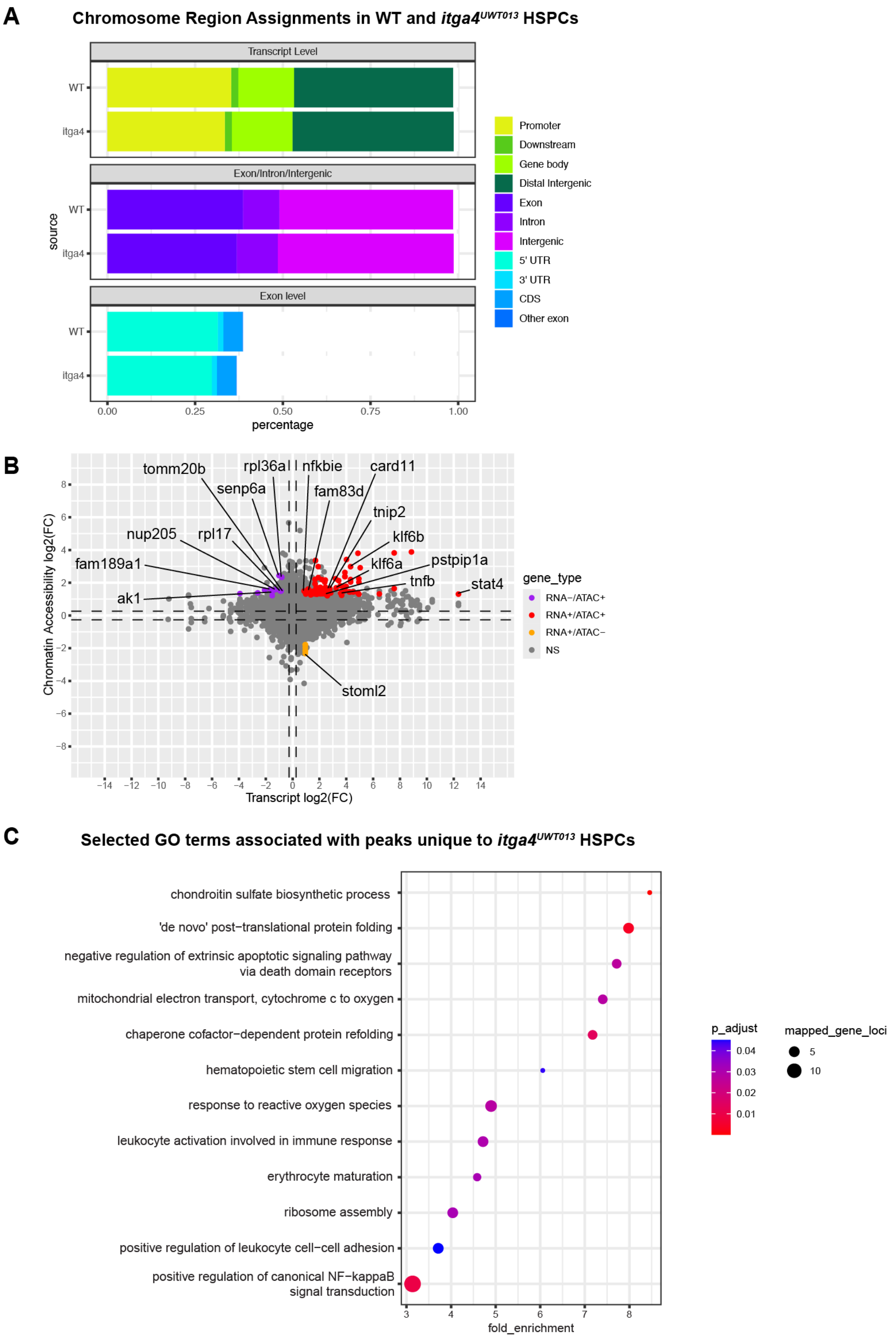


**Supplementary Figure 3. Bulk ATAC-seq of 5 dpf Runx:mCherry+ HSPCs sorted from pooled *itga4*$^{UWT013}$ and WT larvae. Related to Main Figure 3.**

**A)** Charts summarizing the distribution of how called peaks in WT and *itga4*$^{UWT013}$ HSPCs are located within the genome. **B)** Parity plot assessing the correlation of DEGs found from bulk RNA-seq (see **Fig. 1B, Table S1**) with differentially accessible gene loci discovered from bulk ATAC-seq (see **Fig. 3B, Table S3**) in *itga4*$^{UWT013}$ compared to WT HSPCs at 5 dpf. Red dots (top right quadrant) represent gene loci with upregulated accessibility and upregulated transcript expression in *itga4*$^{UWT013}$ compared to WT (denoted as "RNA+/ATAC+" in key); Purple dots (top left quadrant) represent gene loci with upregulated accessibility and downregulated transcript expression in *itga4*$^{UWT013}$ compared to WT (denoted as "RNA-/ATAC+" in key); Orange dots (bottom right quadrant) represent gene loci with downregulated accessibility and upregulated transcript expression in *itga4*$^{UWT013}$ compared to WT (denoted as "RNA+/ATAC-" in key); Grey dots represent gene loci that are not significant (NS), ($p \geq 0.05$); Dashed lines indicate $\log2(FC)=\pm0.25$. Note that the bottom left quadrant is not labeled due to NS data but would be categorized as "RNA-/ATAC-". **C)** Dotplot summarizing the significant gene ontology results of the gene loci with uniquely accessible peaks in *itga4*$^{UWT013}$ HSPCs. Dot size indicates number of DEGs representing GO term, dot color represents significance value (red is low FDR, blue is high), fold enrichment describes *itga4*$^{UWT013}$ compared to WT HSPCs.

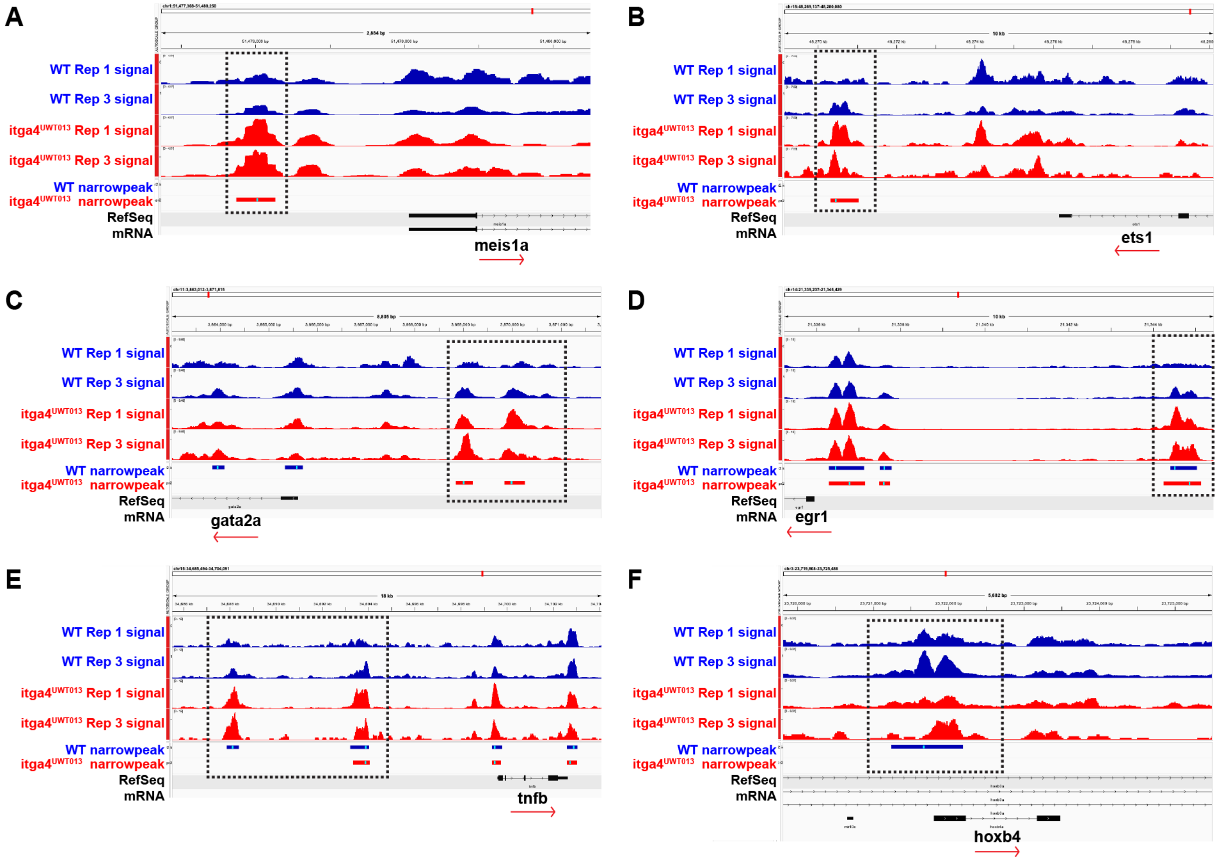


**Supplementary Figure 4. Bulk ATAC-seq tracks at 5 dpf. Related to Main Figure 3.**

**A-F)** Comparison of accessibility tracks between WT (blue upper track) and *itga4*$^{UWT013}$ (red lower track) HSPCs at different gene loci. **A-C)** Genes associated with HSPC identity: **A)** *ets1*; **B)** *gata2a*; **C)** *meis1a*. **D-F)** TFs that have been shown to influence HSPC proliferative activity: **D)** *egr1*; **E)** *tnfb*; **F)** *hoxb4*. Differentially accessible regions, or "peaks", between WT and *itga4*$^{UWT013}$ are indicated by boxes with dashed lines. Track height was auto-scaled by group.

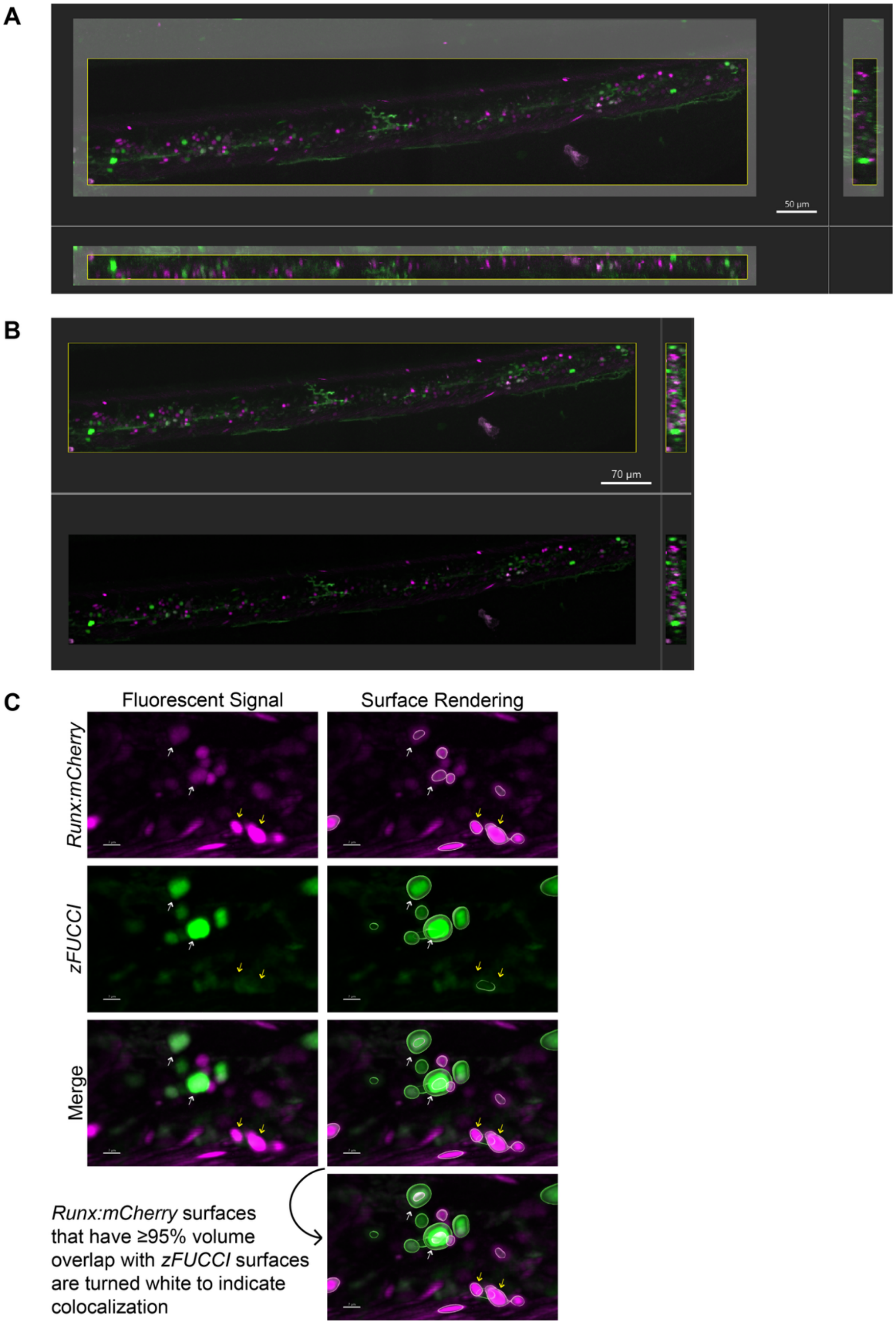


**Supplementary Figure 5. Characterization of the *in vivo* mitotic activity of *itga4*$^{UWT013}$ HSPCs and inflammatory signature of *itga4*-knockdown HSPCs during CHT stage. Related to Main Figure 4.**

**A)** An example of how images were cropped in 3D. Greyed-out areas indicate the sections of the image removed before processing and analysis. Image is of an *itga4*$^{UWT013}$ fish. **B)** An example of the image processing step. The views outlined in the yellow boxes, top left and right, are pre-processed. The views on the bottom are post-processed, with a background subtraction and gaussian filter applied to each channel. Image is of an *itga4*$^{UWT013}$ fish, same as in panel **A**. **C)** An example of rendered surfaces aligning with the transgenic fluorescent signals. White surfaces overlap ≥95% volume with green surfaces. Image is zoomed in on a region from the representative WT fish seen in **Figure 4A**. White arrows indicate the HSPCs with diffuse and/or punctate *Runx:mCherry* expression, expected of cells entering mitosis. Yellow arrows indicate the HSPCs with strong and cohesive *Runx:mCherry* expression, expected of cells in $G_0/G_1$ phases of the cell cycle. Some *Runx:mCherry* expression was too dim to be detected by analysis thresholds (See **Methods** for more details).

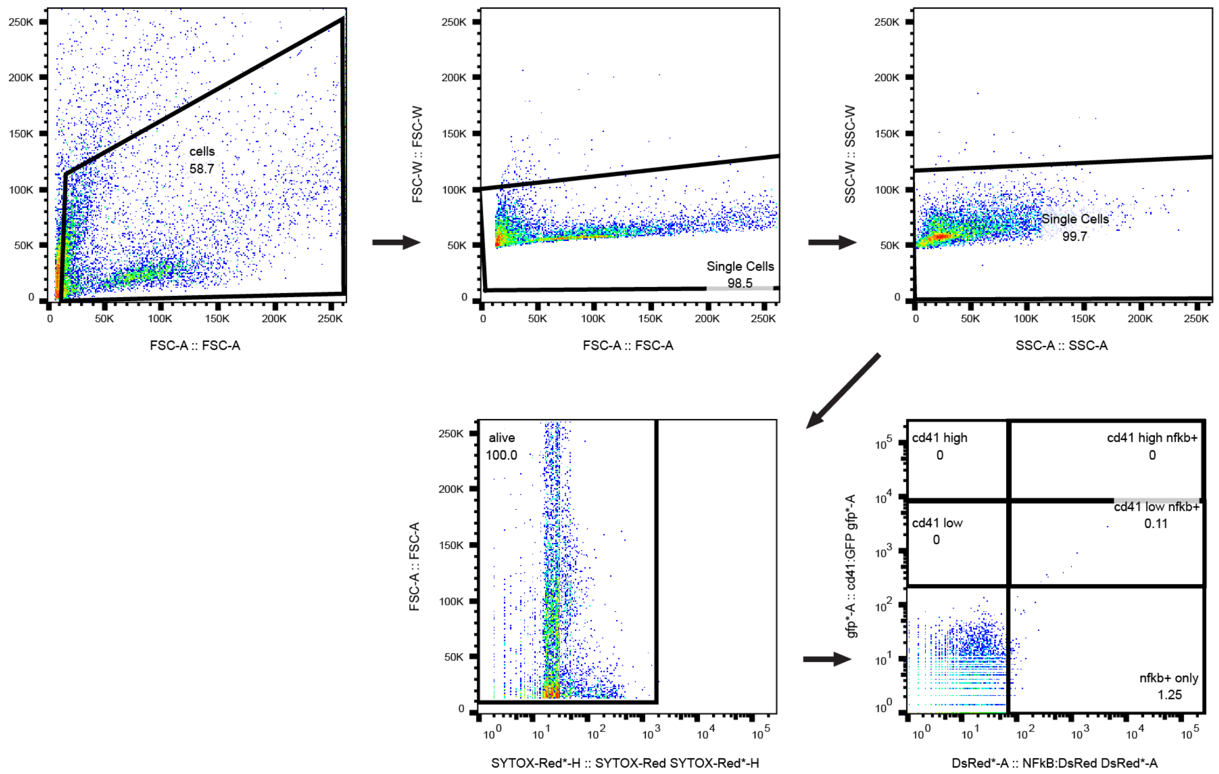


**Supplementary Figure 6. Gating strategy for assessing NF-κB activity in HSPCs during embryonic development via flow cytometry. Related to Main Figure 4.**

Representative gating strategy for flow cytometry showing a representative control sample stained with SYTOX for viability analysis. The final gating step for analysis features cd41:GFP signal intensity on the y-axis and NFκB:d2mRFP signal intensity on the x-axis.

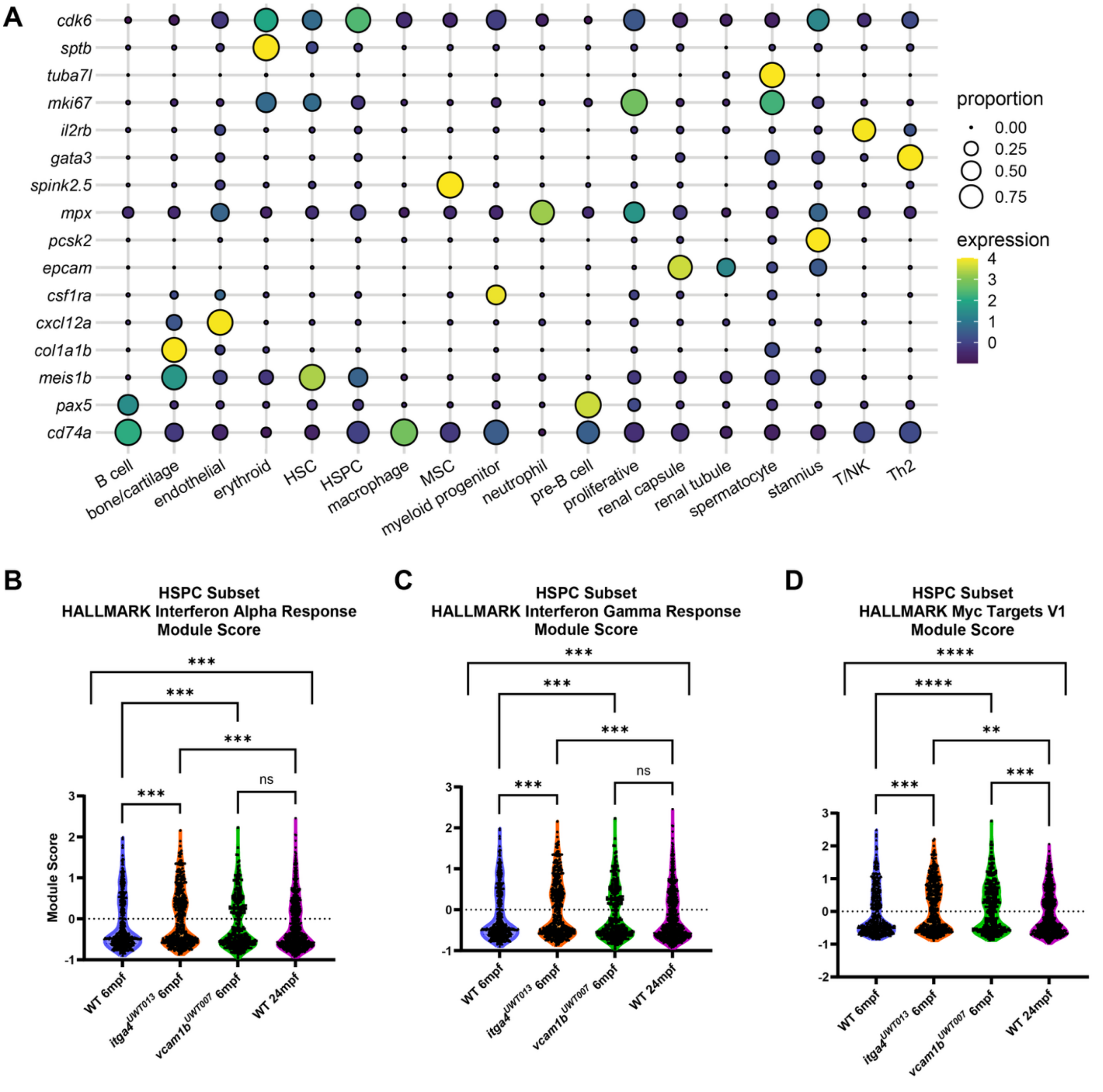


**Supplementary Figure 7. Split-pool barcoding scRNA-seq of adult WKM. Related to Main Figure 5.**

**A)** Bubble plot showing differential expression of individual genes used to determine cluster identities (see **Fig. 5D-E**). **B-D)** Violin Plots comparing module scores assigned to individual cells within the HSPC cluster. A positive module score indicates significantly enriched expression of genes represented by the module dataset, a module score of zero indicates no significant change in module gene set expression compared to control, and a negative module score indicates a significant decrease in the expression of module genes compared to control. Control parameter was set as n=50 genes stochastically selected from each bin. Each symbol inside a violin represents a single cell. The percent of total cells, median, and IQR were calculated independently for the bimodal distribution of each dataset (e.g. HSPCs assigned a negative module score were separated from HSPCs assigned a positive module score within each condition, indicated by horizontal dash line at Module Score=0). Chi-square analysis was performed to compare the proportion of HSPCs assigned negative and positive module scores within each condition, followed by a priori pairwise two-sided Chi-square tests with a Bonferroni correction of $\alpha=0.0125$. Significant chi-square tests and summary statistics reported for each module. In summary statistics, comparison order written as WT 6 mpf vs *itga4* mutant 6mpf vs *vcam1b* mutant 6mpf vs WT 24 mpf (percent indicates proportion of total HSPC population within score category; median and IQR reported for each category). **B)** Chi-square (all conditions)=18.66, degrees of freedom (df)=3, ***p=0.0003; Chi-square (WT 6 mpf vs. *itga4* mutant 6 mpf)=14.32, df=1, ***p=0.0002; Chi-square (*itga4* mutant 6 mpf vs. WT 24 mpf)=14.36, df=1, ***p=0.0002. Positive module score: 31.6% (median=0.610, IQR=0.594) vs 43.6% (median=0.583, IQR=0.628) vs 36.3% (median=0.504, IQR=0.582) vs 32.5% (median=0.472, IQR=0.542); Negative module score: 68.4% (median=-0.535, IQR=0.224) vs 56.4% (median=-0.540, IQR=0.208) vs 63.7% (median=-0.567, IQR=0.218) vs 67.5% (median=-0.575, IQR=0.294). **C)** Chi-square (all conditions)=18.66, df=3, ***p=0.0003; Chi-square (WT 6 mpf vs. *itga4* mutant 6 mpf)=14.32, df=1, ***p=0.0002; Chi-square (*itga4* mutant 6 mpf vs. WT 24 mpf)=14.36, df=1, ***p=0.0002. Positive module score: 31.6% (median=0.610, IQR=0.594) vs 43.6% (median=0.583, IQR=0.628) vs 36.3% (median=0.504, IQR=0.582) vs 32.5% (median=0.472, IQR=0.542); Negative module score: 68.4% (median=-0.535, IQR=0.224) vs 56.4% (median=-0.540, IQR=0.208) vs 63.7% (median=-0.567, IQR=0.218) vs 67.5% (median=-0.575, IQR=0.294). **D)** Chi-square (all conditions)=25.77, df=3, ****p<0.0001; Chi-square (WT 6 mpf vs. *itga4* mutant 6 mpf)=14.5, df=1,

***p=0.0001; Chi-square (WT 6 mpf vs. *vcam1b* mutant 6 mpf)=16.86, df=1, ****p<0.0001; Chi-square (*itga4* mutant 6 mpf vs. WT 24 mpf)= 8.723, df=1, **p=0.0031; Chi-square (*vcam1b* mutant 6 mpf vs. WT 24 mpf)=11.02, df= 1, ***p=0.0009. Positive module score: 35.9% (median=0.634, IQR=0.720) vs 48.3% (median=0.752, IQR=0.637) vs 50.3% (median=0.641, IQR=0.664) vs 39.4% (median=0.592, IQR=0.633); Negative module score: 64.1% (median=-0.537, IQR=0.246) vs 51.7% (median=-0.549, IQR=0.232) vs 49.7% (median=-0.549, IQR=0.246) vs 60.6% (median=-0.575, IQR=0.293). Modules were created from HALLMARK Human Gene Sets: **B)** HALLMARK Interferon Alpha Response; **C)** HALLMARK Interferon Gamma Response; **D)** HALLMARK Myc Targets V1.

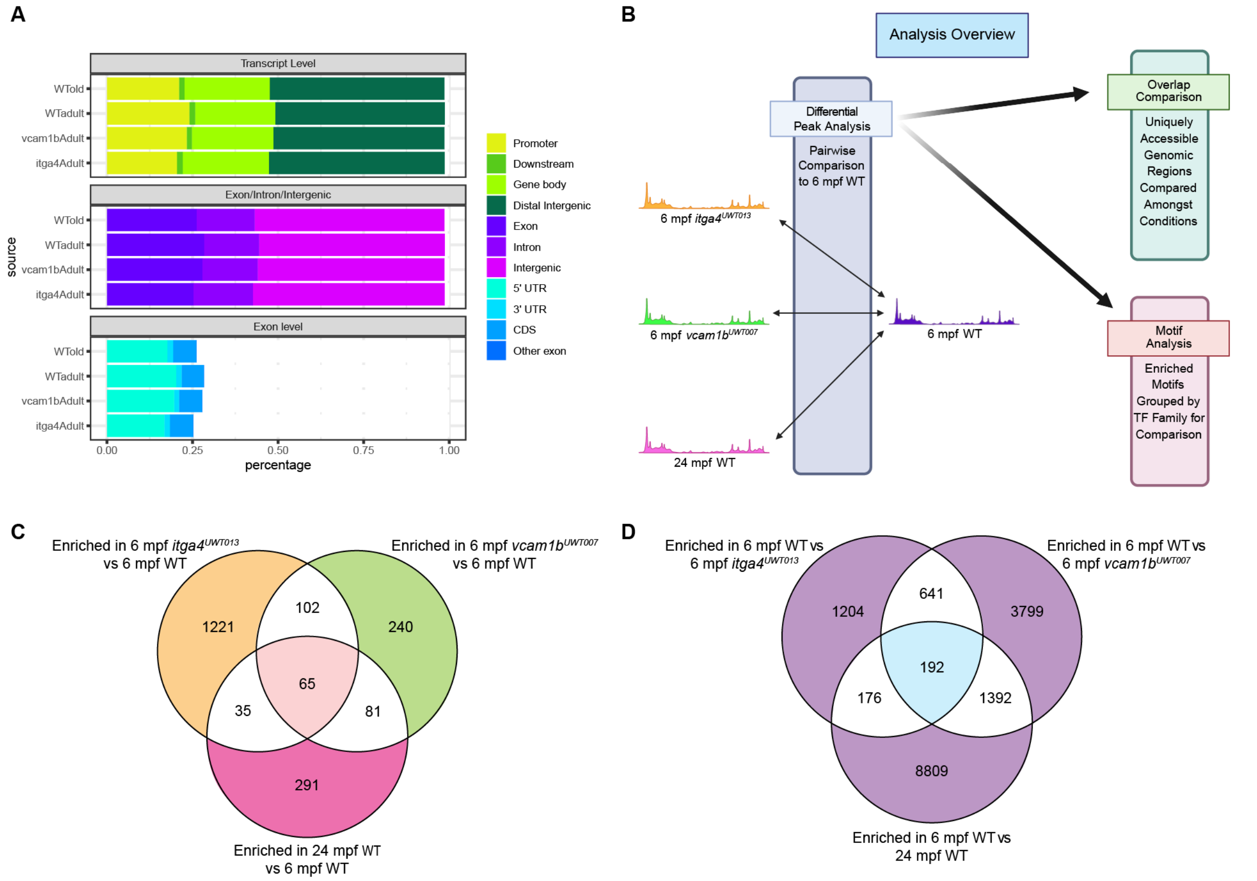


**Supplementary Figure 8. Bulk ATAC-seq of Runx:mCherry+ HSPCs sorted from Adult WKM. Related to Main Figure 6.**

**A)** Charts summarizing the distribution of how called peaks in adult Runx:mCherry+ HSPCs from each condition are located within the genome. **B)** Schematic describing the steps taken during the analysis stage. Young adult (6 mpf) WT was treated as the "control" by which all other conditions were compared for diffbind, HOMER, and TOBIAS analyses. **C-D)** Venn diagrams representing the results of differential peak analysis via pairwise comparison to 6 mpf WT for each condition (**C**: orange, green, or pink outer edges of circles indicates enrichment in 6 mpf *itga4*$^{UWT013}$, 6 mpf *vcam1b*$^{UWT007}$, or 24 mpf WT respectively; **D**: purple outer edges of circles indicates enrichment in 6 mpf WT) and the results of overlap comparisons to identify accessibility in shared genomic regions. **C)** The number of genes featuring enriched accessibility from each comparison to 6 mpf WT. Enriched gene loci shared by 6 mpf *itga4*$^{UWT013}$, 6 mpf *vcam1b*$^{UWT007}$, and 24 mpf WT HSPCs compared to 6 mpf WT HSPCs colored in red, center panel. **D)** The number of genes featuring decreased accessibility from each comparison to 6 mpf WT. Decreased gene accessibility shared by 6 mpf *itga4*$^{UWT013}$, 6 mpf *vcam1b*$^{UWT007}$, and 24 mpf WT HSPCs compared to 6 mpf WT HSPCs colored in blue, center panel.

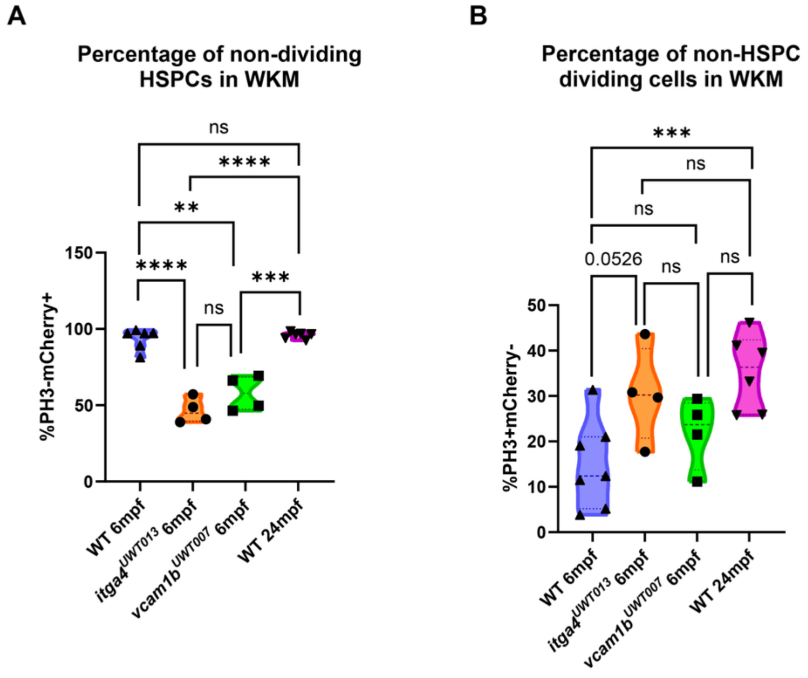


**Supplementary Figure 9. Adult WKM Flow. Related to Main Figure 7.**

**A-B)** Percentages were analyzed using generalized linear models for overdispersed binomial data. P-values for comparisons among groups were adjusted for multiplicity using the single-step procedure described in Hothorn et al.[1] that relies on the multivariate t-distribution for all pairwise comparisons. **A)** Violin plot of non-proliferating HSPCs (negative for staining by conjugated AlexaFluor647:PH3 antibody) given as a percentage of the total Runx:mCherry+ population in WKM. Parent gate is live Runx:mCherry+ single cells (not shown). The purple violin with triangle points represents 6 mpf WT (n=6 biological replicates; median=97.2; IQR=10.6), the orange violin with circle points represents 6 mpf *itga4*$^{UWT013}$ (n=4 biological replicates; median=44.9; IQR=15.7), the green violin with square points represents 6 mpf *vcam1b*$^{UWT007}$ (n=4 biological replicates; median=57.9; IQR=21.3), and the pink violin with upside-down triangle points represents 24 mpf WT (n=6 biological replicates; median=96.5; IQR=3.72). One WKM from one adult fish was defined as one biological replicate, and sample processing occurred on three independent experiment days. Significant comparisons are as follows: 6 mpf WT vs 6 mpf *itga4*$^{UWT013}$

(****p<0.0001); 6 mpf WT vs 6 mpf *vcam1b*$^{UWT007}$ (**p=0.0018); 24 mpf WT vs 6 mpf *itga4*$^{UWT013}$ (****p<0.0001); 24 mpf WT vs 6 mpf *vcam1b*$^{UWT007}$ (***p=0.0007). **B)** Violin plot of proliferating cells that are not HSPCs (Runx:mCherry-) given as a percentage of the total PH3+ population in KM. Parent gate is all live single cells excluding red blood cell lysis (not shown). The purple violin with triangle points represents 6 mpf WT (n=7 biological replicates; median=12.4; IQR=15.8), the orange violin with circle points represents 6 mpf *itga4*$^{UWT013}$ (n=4 biological replicates; median=30.2; IQR=19.7), the green violin with square points represents 6 mpf *vcam1b*$^{UWT007}$ (n=4 biological replicates; median=23.7; IQR= 4.8), and the pink violin with upside-down triangle points represents 24 mpf WT (n=6 biological replicates; median=36.4; IQR=16.5). One WKM from one adult fish was defined as one biological replicate, and sample processing occurred on three independent experiment days. Significant comparisons are as follows: 6 mpf WT vs 24 mpf WT (***p=0.0008).

1. Hothorn T, Bretz F, Westfall P. Simultaneous inference in general parametric models. *Biom J* **50**, 346-363 (2008).
